# *Ggnbp2* regulates synaptic development and autophagy in motor neurons

**DOI:** 10.1101/2023.11.03.565470

**Authors:** Sarah K. Kerwin, Nissa Carrodus, Amber Kewin, Tian Lin, Xiaoyu Qian, Allan F. McRae, Jian Yang, Brett M. Collins, Naomi R. Wray, Fleur C. Garton, S. Sean Millard

**Affiliations:** School of Biomedical Sciences, Faculty of Medicine, The University of Queensland, Brisbane, 4072, Australia; Institute for Molecular Biosciences, The University of Queensland, Brisbane, 4072, Australia; School of Life Sciences, Westlake University, Hangzhou, 310024, China; Westlake Laboratory of Life Sciences and Biomedicine, Hangzhou, 310024, China; Queensland Brain Institute, The University of Queensland, Brisbane, 4072, Australia

## Abstract

Genome-wide association studies (GWAS) have identified numerous candidate ALS risk variants, but their cellular functions are often unknown. Recent studies have identified a variant of *GGNBP2* that results in increased expression. To better understand how this gene might contribute to disease, we investigated the function of *Drosophila Ggnbp2 (dGgnbp2)* in motor neurons. Loss of function studies showed that *dGgnbp2* is required for motor neuron synaptic development. A human transgene completely rescued these phenotypes indicating that the gene is functionally conserved between humans and flies. Overexpression of *dGgnbp2* caused severe locomotor defects in adult flies, consistent with ALS pathology. At the cellular level, *dGgnbp2* regulated autophagy, a process commonly defective in ALS. Both overexpression and removal of *dGgnbp2* reduced levels of the phosphorylated lipid, PI(3)P, an essential component of autophagosomes. Our study provides strong evidence that *Ggnbp2* functions in motor neurons to regulate a cellular process commonly defective in ALS.

**Teaser:** This study investigated the function of the ALS risk variant *GGNBP2*, in flies, and showed that it regulates autophagy in motor neurons.

## Introduction

The gene *Gametogenetin Binding Protein 2* (*GGNBP2*) was associated with ALS at the level of genome-wide significance using a gene-based test, which accumulates association signal from SNPs across the gene (*1*). In the same study, the SNP associations were integrated with gene expression (eQTL) data using the summary data-based Mendelian randomization (SMR) method, for which the most significant test result again implicated *GGNBP2* (*1*). The human ALS associated locus is a region of high linkage disequilibrium with SNPs in neighboring genes that are highly correlated. Here we carried out follow-up studies using larger datasets and hypothesized that functional studies would demonstrate that *GGNBP2* plays a role in motor neurons and that overexpression of the protein (associated with risk allele) would perturb its normal function.

*GGNBP2* was first identified as a gene induced by the environmental pollutant, dioxin, in mouse embryonic stem cells. Given its high level of expression in mouse testis, it was suggested that dioxin’s reproductive toxicity could be related to *GGNBP2* misexpression (*2*). *GGNBP2* was later identified in a yeast 2-hybrid screen using the germ cell specific protein, gametogenetin (GGN), as bait (*3*). Further research demonstrated that GGNBP2 plays a developmental role, potentially as a C2H2 Zn-finger transcription factor, in both placenta and testis (*4-6*). Cancer studies have indicated that GGNBP2 may act as a tumor suppressor able to inhibit several growth-promoting signal transduction pathways (*7-9*). The gene is expressed in most tissues (with highest levels in testis) (*10*), and its cellular localization and function may be cell-specific as the protein has been detected in both the nucleus and the cytoplasm (*2, 3*). A role for *GGNBP2* in neurons has not been previously described.

Given that 75% of genes involved in human disease are conserved in *Drosophila melanogaster* (*11*), we exploited this system to validate *GGNBP2* as an ALS risk factor and to better understand its function related to this disease. We used CRISPR to delete the fly *dGgnbp2* homolog and found significant changes in motor neuron morphology and synaptic makeup. These phenotypes were completely rescued by expressing human *GGNBP2* in motor neurons, demonstrating functional conservation and an autonomous requirement in the type of neuron most affected in ALS. Overexpression of the gene caused motor neuron phenotypes opposite to that of the null mutant and a severe locomotion defect in adult flies. dGgnbp2 protein localized primarily to the cytoplasm of motor neurons and we confirmed that it functions in the cytoplasm by rescuing mutant phenotypes with a *dGgnbp2* transgene lacking a nuclear localization signal. RNA-seq on *dGgnbp2* mutant flies showed a downregulation of autophagy, endocytosis and phosphatidylinositol signaling. Consistent with this, we found that *dGgnbp2* genetically interacts with another ALS risk factor, *TANK-binding kinase 1 (TBK1)* (*IKK-related kinase*, *ik2* in flies) that plays a role in autophagy. Both *dGgnbp2* overexpression and loss of function delayed autophagic flux. Its role in autophagy involved PI3K regulation as *dGgnbp2* overexpression promoted class I kinase activity, which inhibits autophagy, and reduced lipid species generated by class III PI3K that are critical for autophagosome maturation.

## Results

### *GGNBP2* is the top chromosome 17 ALS candidate risk gene using the latest GWAS results

*GGNBP2* was originally identified as a putative risk gene in a cross-ancestry GWAS of 13,811 sporadic ALS cases and 26,350 controls (*1*). Here, we use the latest (and largest) ALS GWAS results based on 29,612 sporadic cases and 122,656 controls (Table 1) (*12*). The SNP most significantly associated with ALS in this region (rs11650008, p=8.6×10^-8^) is annotated as an intronic variant of a neighboring gene, *PIGW*. GCTA-COJO analysis (*13*) (used to check for secondary associations in the region) provided support for only a single association (Table S1). Gene-based tests (*14*) applied to the GWAS data (*12*) identified *GGNBP2* as the most significantly associated gene in the region (Table S2), retaining the genome-wide significance previously reported (p=6.6 x10^-7^) (Figure 1A, Table S3). We note that four other genes in the region also pass the genome-wide significance threshold (*DHRS11*, *PIGW*, *MYO19* and *ZNHIT3*) as their gene boundaries share SNPs that are highly correlated (Figure 1A-B, Table S4). None of these genes were enriched for rare variants from the Project MinE study in 4,633 cases vs.1,832 controls (Table S5). To determine whether the ALS-associated SNP correlated with changes in gene expression in any of the five genes in this region, gene expression data (eQTL) from human blood were integrated with GWAS results using the summary data-based Mendelian randomization (SMR) method. *GGNBP2* had the largest change in gene expression compared to the non-risk allele and the lowest SMR *p* value (Figure 1A-B). We replicated this result using a smaller (but largest available) brain eQTL dataset (Table S5). Plotting the effect size of expression (eQTL) data against the GWAS effect size clearly implicated *GGNBP2* as the top gene in this region (Figure 1C). In summary, these analyses based on the latest available data prioritise *GGNBP2* within this chromosome 17 risk locus.

**Figure 1.**
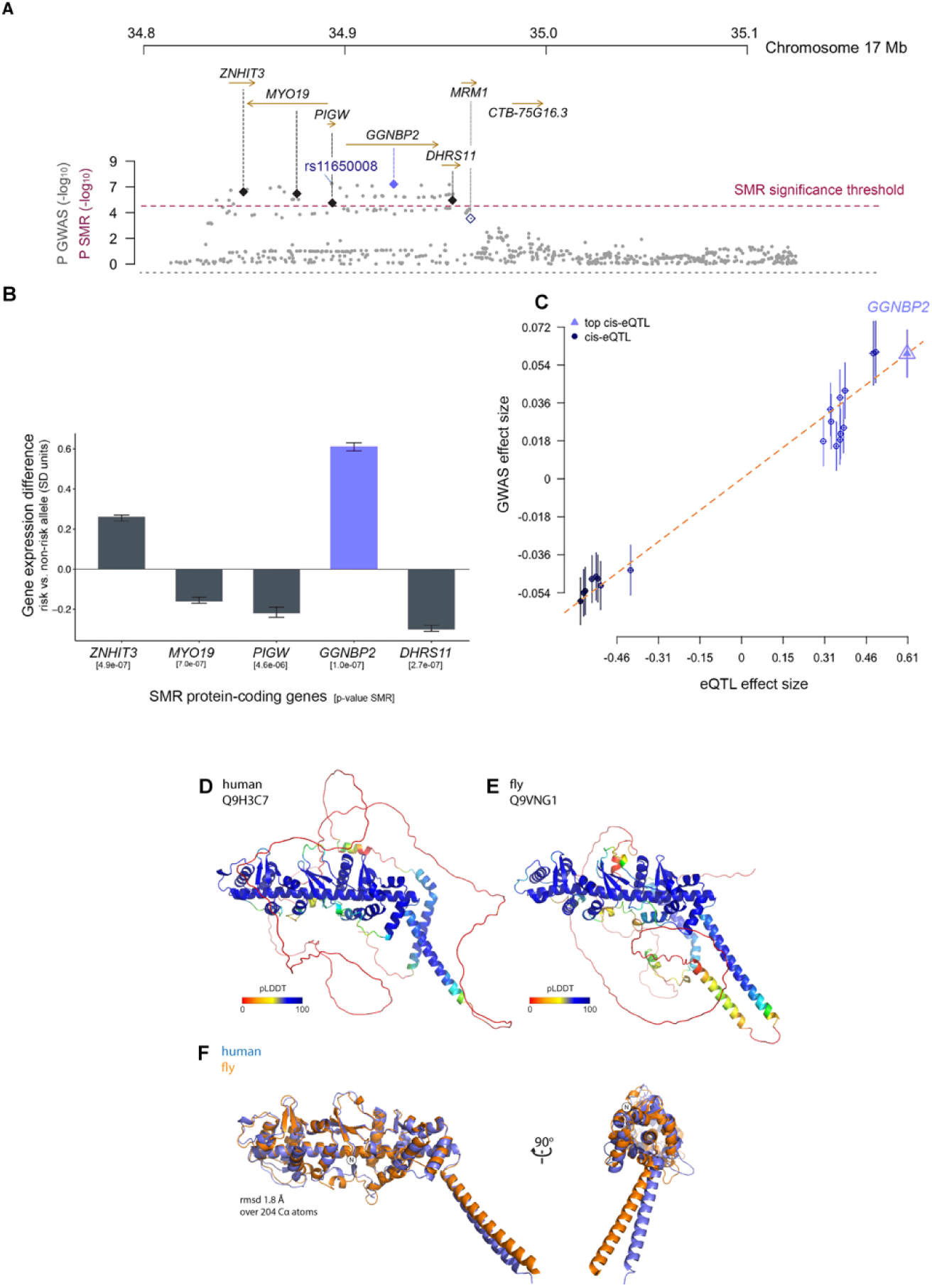
GWAS post-analyses identify *GGNBP2* as an ALS candidate risk gene. (A)The Summary-based Mendelian Randomisation (SMR) test results for the locus on Chromosome 17 (which quantifies the statistical evidence of the ALS associated SNP (ALS GWAS data (*12*)) being mediated through the SNP control of gene expression (eQTL data, (*62*)). The position and direction of the region’s genes are shown, with five whose SNP ALS association is genome-wide significant (using the SMR threshold (as labelled)) (*p* <8.4 e-6). Plotted on the y-axis are both -log_10_ *p* values for ALS-associated SNPs (grey dots) and for the SMR test which integrates the eQTL data with the GWAS results (diamonds). The top GWAS SNP (rs11650008) is in blue. The x-axis corresponds to genomic location. The linkage disequilibrium (correlation between SNPs) is evident from the block of high -log_10_ *p* values consistent across the SMR plot. This includes every SNP available in the region in the GWAS and eQTL summary data. *GGNBP2* has the lowest *p* value for both the SMR data and the gene-based GWAS test. (B) The gene expression difference associated with the top ALS GWAS- SNP in human blood (N=31,684, eQTL data, (*62*)). On the y axis is the ratio of the risk allele compared to the non-risk allele (expressed in standard deviation units, error bars 95% confidence intervals). The x-axis includes the gene and its SMR *p* value (from A). (C) Effect sizes of SNPs from GWAS (y-axis) plotted against effect sizes for SNPs from the eQTL dataset (x-axis). The top cis-eQTL (triangle) correlates to *GGNBP2*. The SNPs are shaded according to their r^2^ with the top-SNP (1=royal blue and 0=black). (D-E) Predicted structure of human GGNBP2 (D) and *Drosophila melanogaster* dGgnbp2 (E) proteins by AlphaFold2. Structures are colored based on the pLDDT confidence score (*17, 75*). (F) Overlay of the N-terminal SRD of GGNBP2 of human (blue) and fly (orange) proteins predicted by AlphaFold2 shows excellent structural correlation. See also Figure S1 and S2.

**Table 1.** Top gene for each chromosome using gene-based test (mBAT-combo) ordered by p-value. (see Table S2 for all results). The topSNP is based on the ALS GWAS European results from van Rheenen et al. 2021.

| Gene | Chr | Start | End | No.SN<br>Ps | TopSNP | TopSN<br>P_Pval<br>ue | No.Eig<br>enval<br>ues | Chisq<br>_mBA<br>T | P_mBAT<br>combo |
| --- | --- | --- | --- | --- | --- | --- | --- | --- | --- |
| **UNC13A** | 19 | 17712145 | 17799174 | 92 | rs12608932 | 6.63E-25 | 11 | 108.0 | 9.23E-18 |
| **SLC9A8** | 20 | 48429250 | 48508779 | 229 | rs17785991 | 3.99E-10 | 6 | 49.6 | 3.53E-10 |
| **CFAP410** | 21 | 45748827 | 45759285 | 15 | rs75087725 | 2.65E-11 | 6 | 49.8 | 1.04E-08 |
| **GGNBP2** | 17 | 34900761 | 34946278 | 60 | rs9903355 | 1.30E-07 | 4 | 29.7 | 6.63E-07 |
| **FSD2** | 15 | 83424114 | 83474822 | 106 | rs2379994 | 1.45E-07 | 8 | 40.2 | 9.37E-07 |
| **MEF2C** | 5 | 88012934 | 88200074 | 302 | rs17558396 | 6.01E-07 | 16 | 46.4 | 1.26E-06 |
| **KIF5A** | 12 | 57939809 | 57980416 | 55 | rs113247976 | 1.46E-11 | 11 | 43.8 | 1.43E-06 |
| **LY6G5C** | 6 | 31644461 | 31651817 | 6 | rs805290 | 2.10E-06 | 4 | 31.5 | 1.67E-06 |
| **ZSWIM8** | 10 | 75545364 | 75561555 | 6 | rs2271271 | 5.20E-05 | 3 | 30.7 | 1.94E-06 |
| **PROS1** | 3 | 93591895 | 93698847 | 74 | rs13089004 | 6.83E-07 | 9 | 35.1 | 2.00E-06 |
| **CLCN3** | 4 | 170533784 | 170644824 | 224 | rs62333164 | 6.29E-08 | 6 | 29.3 | 2.04E-06 |
| **FNBP1** | 9 | 132649466 | 132805473 | 456 | rs7870884 | 1.92E-06 | 16 | 35.9 | 3.31E-06 |
| **SFI1** | 22 | 31884674 | 32014574 | 382 | rs55894054 | 3.85E-05 | 6 | 35.7 | 4.97E-06 |
| **WIPI2** | 7 | 5229827 | 5273471 | 140 | rs6961379 | 0.00014 | 8 | 40.3 | 5.51E-06 |
| **PSMB10** | 16 | 67968409 | 67970767 | 2 | rs14178 | 2.02E-06 | 2 | 23.1 | 8.16E-06 |
| **RESP18** | 2 | 220192129 | 220197899 | 17 | rs6710951 | 7.09E-07 | 3 | 25.6 | 1.06E-05 |
| **ILK** | 11 | 6624938 | 6632105 | 28 | rs79548722 | 5.34E-06 | 3 | 26.6 | 1.07E-05 |
| **TRIP11** | 14 | 92432335 | 92507240 | 148 | rs2295170 | 5.55E-05 | 8 | 36.5 | 2.80E-05 |
| **NVL** | 1 | 224415036 | 224517891 | 183 | rs4653570 | 5.54E-07 | 22 | 47.8 | 4.69E-05 |
| **COG3** | 13 | 46039033 | 46110836 | 179 | rs3014966 | 2.49E-06 | 5 | 23.1 | 7.14E-05 |
| **C8orf34** | 8 | 69243190 | 69731258 | 1309 | rs10099635 | 0.0020 | 48 | 94.1 | 0.00016 |
| **GRP** | 18 | 56887390 | 56898006 | 38 | rs34973755 | 0.0019 | 4 | 18.1 | 0.0018 |

### *dGgnbp2* is required in motor neurons

The fly orthologue of *GGNBP2*, *CG2182* (hereafter referred to as *dGgnbp2*), encodes a protein of 671 amino acids that is 29% identical (43% similar) to the 697 amino acid human protein (Figure S1) (*15, 16*). AlphaFold2 predicts the structures of the human and fly proteins are very similar (Figure 1C-E, S2) (*17, 18*). GGNBP2 has an N-terminal domain predicted to have a previously undescribed fold and a disordered C-terminal domain. The N-terminal domain begins with a short β-sheet (βD) and consists of two repeated structures we call D1 and D2 that wrap around a long central α-helix (H1) (Figure 1E). This architecture is reminiscent of a ‘sausage-roll’ and appears to be unique to GGNBP2 homologues based on database searches with FoldSeek; we propose to name it the SRD (<u>S</u>ausage <u>R</u>oll domain). Notably the SRD was found to be necessary and sufficient for interacting with its known binding partner, gametogenetin, using Alphafold2. The C-terminal domain of GGNBP2 has recently been shown to be necessary for binding to the CCR4-NOT deadenylase protein complex (*19*). Whether either of these interactions occur in neurons is not known.

To interrogate the function of this gene, we generated a null allele using CRISPR (*dGgnbp2^null^*), by deleting the entire coding region (Figure 2A). Homozygous null animals were viable and fertile. To assess whether motor neurons were perturbed in *dGgnbp2^null^* animals, we analyzed morphologic and synaptic phenotypes of the well characterized larval motor neuron that innervates muscle 6 and 7 (MN6/71b). This glutamatergic neuron forms round synaptic boutons on postsynaptic muscle, each bouton containing many active zones that can be visualized with the synaptic scaffolding protein brp (*20*). Compared to heterozygotes (Figure S3A-F) and controls, *dGgnpb2^null^* motor neurons showed a marked reduction in bouton size (surface area in μm^2^), an increase in bouton number (normalised to muscle size) and a reduction in synapses (brp puncta/bouton), (Figure 2B-F). The increase in bouton number translated to a net increase in the number of brp puncta per neuromuscular junction (NMJ) (Figure 2G). Muscle size (surface area in μm^2^) was also reduced in *dGgnpb2^null^* (Figure S3G). As a readout for the microtubule network required for stable synapses, we used the microtubule binding protein Futsch, which labels microtubules in axon shafts and forms circular structures in a subset of boutons (*21*). Futsch is a translational target of TDP-43, a protein that forms insoluble cytoplasmic aggregates in ALS and other neurodegenerative diseases (*22*). The expression and localisation of Futsch is frequently perturbed in fly models of ALS (*23*)*. dGgnpb2^null^* (not *dGgnpb2^null/+^* (Figure S3H-K)) animals showed reduced and diffuse Futsch labelling, as well as a decrease in loops (mature representations of microtubules) (Figure 2H-L). In contrast, no discernible differences in glutamate receptor (GluRIIA) density or intensity were observed in the muscle of *dGgnbp2^null^* animals compared to controls (Figure S3L-O).

**Figure 2.**
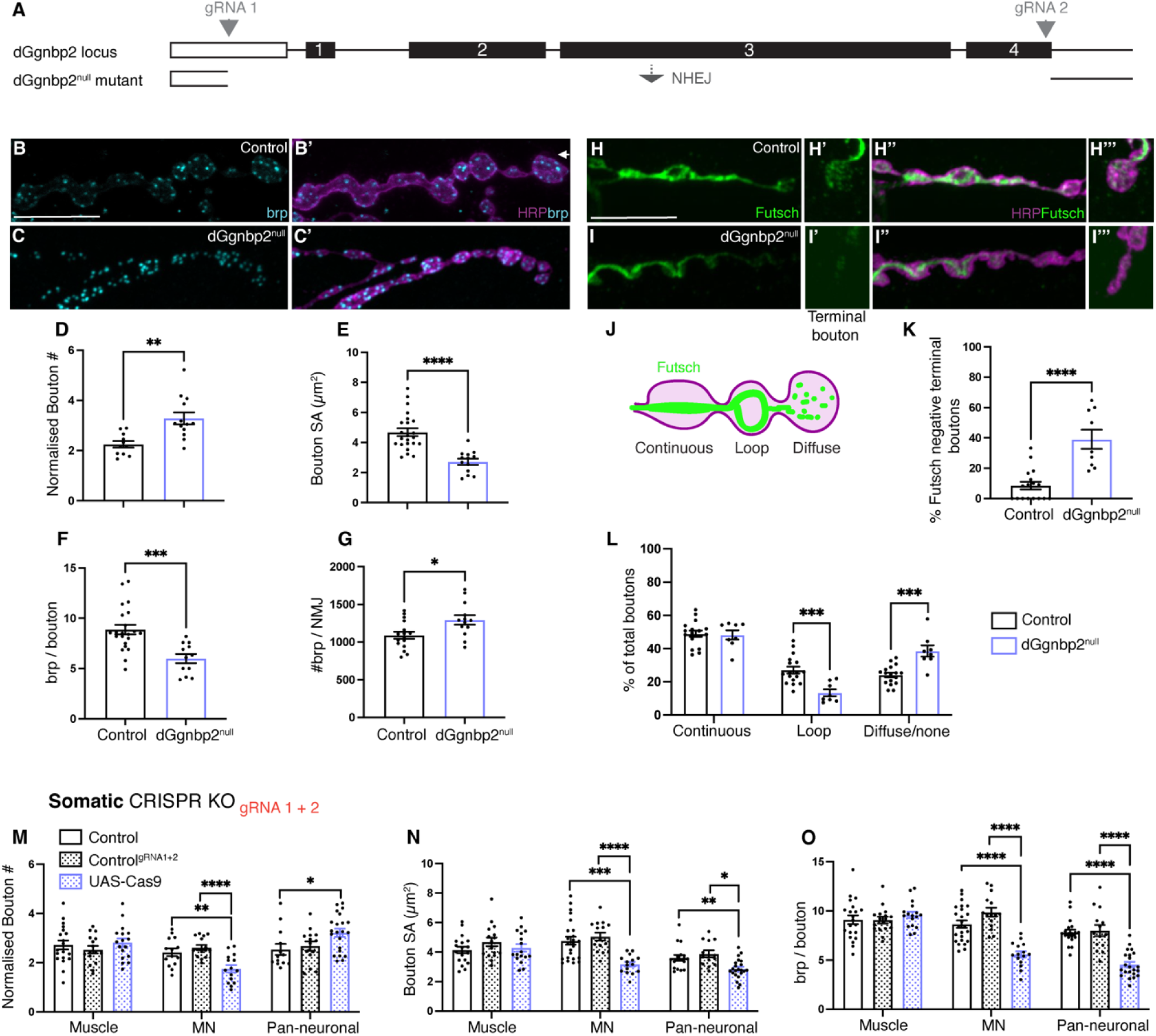
*dGgnbp2* is required in motor neurons. (A) Schematic of *dGgnbp2^null^* CRISPR mutant. (B-B’) Control (*w^1118^*) MN6/71b labelled with nc82 antibody (cyan, B) to mark brp positive active zones and merged with HRP (magenta, B’) which labels the axonal membrane. (C-C’) *dGgnbp2^null^*MN6/7 1b with the same labelling scheme as B. (D-G) Quantification of bouton number normalised to muscle 6/7 surface area (1 x 10^3^μm^2^) (D) bouton size (E) brp puncta/bouton (F) and the number of brp puncta/neuromuscular junction (G). (H-I) Futsch labelling of Control (*w^1118^*) MN6/71b labelled with antibody to futsch (green, H, H’) and merged with HRP (magenta, H’’, H’’’). High magnification of a terminal bouton showing a futsch loop (H’, H’’’). *dGgnbp2^null^* motor neuron with the same labelling scheme as in H (I-I’’’). (J) Schematic of the microtubule binding protein, futsch, in boutons. Continuous futsch tracts and loops are indicative of stable synapses. (K-L) Quantification of futsch in terminal boutons (K) and the different states of futsch labelling in control and *dGgnbp2^null^* boutons. n=8-16 (L). (M-O) Quantification of MN phenotypes as in D-F using somatic CRISPR to knockout *dGgnbp2* in muscle and motor neurons. Control (*Gal4* alone: *BG487* (muscle) and *OK6* (MNs)) (black outline), Control (*Gal4,* gRNAs) (black stippled) and *dGgnbp2* somatic knockout (*Gal4*, gRNAs and *UAS-Cas9*) (purple stippled). n=12-24, D-G, K-O. All quantitative analyses were performed on compressed Z stacks of spinning disk microscopy and analysed using ImageJ. Each data point=average of 10 boutons. Data are represented as mean ± SEM. Depending on distribution of data an unpaired t test (D-G) or Mann-Whitney test (K-L) or an Ordinary one-way ANOVA with Tukey’s multiple comparison (M^OK6, BG487^, O^OK6, BG487,elav^), test or Kruskal-Wallis test with Dunn’s multiple comparison test (M^elav^, N^OK6, BG487,elav^) was used. n.s *p*>0.05, \**p*<0.05, \*\*\**p*<0.001 and \*\*\*\**p*<0.0001. White arrows show MN6/71b. Scale bars denote 10um. See also Figure S3.

To confirm that these phenotypes were due to a loss of *dGgnpb2* and not an off-target effect from CRISPR, we looked at transheterozygotes of *dGgnbp2^null^* and chromosomal deletion lines (*Df(3R)BSC464* and *Df(3R)BSC47*) (*24*). These lines contain a deletion that encompasses the *dGgnbp2* locus. The reductions in brp puncta and bouton size observed in d*Ggnbp2^null^* animals were replicated in these transheterozygotes (Figure S3P-Q).

To determine whether *dGgnbp2* is autonomously required in motor neurons, we used somatic CRISPR to drive expression of *Cas9* in specific tissues (muscle, motor neurons, pan-neuronal) while constitutively expressing two *dGgnbp2 gRNAs*. As a proof of principle that somatic CRISPR is effective at removing *dGgnbp2*, we targeted *Cas9* to all neurons and recapitulated the *dGgnbp2^null^* phenotypes. Removing *dGgnbp2* from muscle did not affect motor neuron morphology or synaptic makeup, whereas removing it specifically from motor neurons resulted in similar motor neuron defects as observed in the *dGgnbp2^null^* animals (Figure 2M-O). A *dGgnbp2* requirement in motor neurons was confirmed by rescuing the *Ggnbp2^null^* phenotypes with a V5-tagged version of *dGgnbp2* in motor neurons. Driving this same transgene in muscle failed to rescue (Figure S3R-X). Together, these data demonstrate that *dGgnbp2* functions in a cell autonomous manner to support motor neuron morphology and synaptic development.

### Overexpressing *Ggnbp2* in motor neurons induces axon morphology and synaptic composition deficits

Using large eQTL datasets (not enriched for disease) individuals carrying the *GGNBP2* risk allele exhibit increased expression of this gene in both brain and blood (Figure 1B, Figure S1, Table S6). Therefore, we assessed the effects of *dGgnbp2* overexpression at the *Drosophila* NMJ. To achieve overexpression we utilised both a *UAS P-element* insertion in the 5’ end of the gene (*EPgy2*), allowing for *Gal4*-dependent expression of *dGgnbp2*, hereafter referred to as *dGgnbp2^OE-EPgy2^* (Figure 3A) (*25, 26*) and a V5-tagged UAS transgene (*dGgnbp2*^V5^). We drove *dGgnbp2* expression in motor neurons using *OK6-Gal4* and assessed motor neuron morphology, synaptic composition, and the cytoskeletal organisation of the neuron MN6/71b, similar to what was done for the *dGgnbp2^null^* allele above. Notably, overexpression phenotypes had an inverse relationship with the null allele effects. Motor neuron bouton size (surface area in μm^2^) was increased in *dGgnbp2^OE-EPgy2^* with a corresponding decrease in the number of boutons (normalised to muscle size) (Figure 3B-G, S4A-H). The number of brp puncta (synapses) were also increased per bouton however, a net decrease in synapses was observed across the NMJ (Figure 3G). Muscle size (surface area in μm^2^) was also reduced in *dGgnpb2^OE-EPgy2^* when overexpression was restricted to motor neurons (Figure S4I-J). Diffuse or no Futsch labelling was observed when *dGgnbp2* was overexpressed (*dGgnbp2^OE-EPgy2^*), particularly in terminal boutons (Figure 3H-K). In contrast to these presynaptic defects, glutamate receptor (GluRIIA) density remained unchanged, although a slight increase in fluorescence was observed (Figure S4K-N).

**Figure 3.**
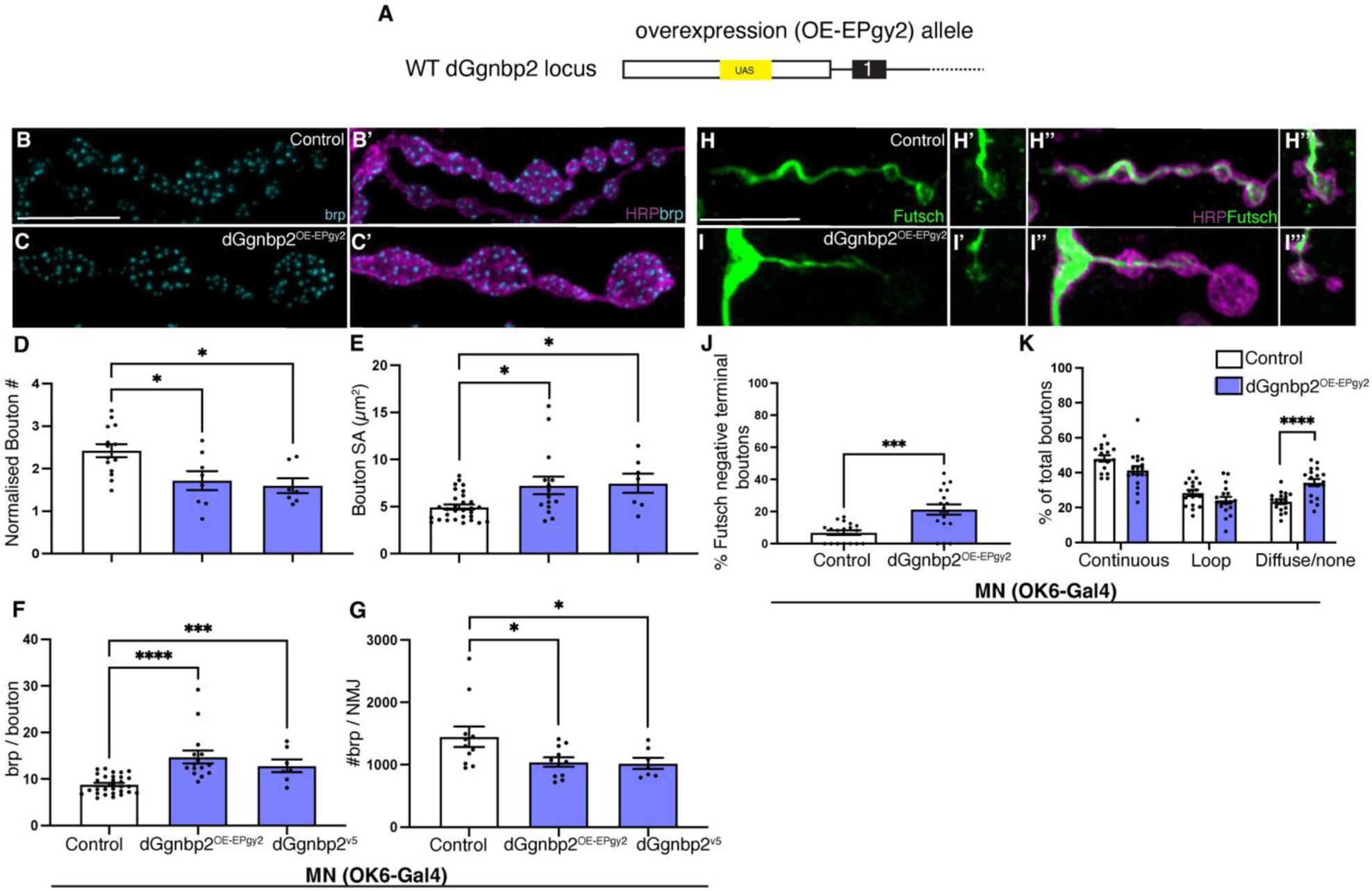
Overexpressing *dGgnbp2* in motor neurons changes axon morphology and synaptic composition. (A) Schematic showing the *P*[*EPgy2*](EY00712) (*25*) insertion in the 5’ region of endogenous *dGgnbp2* used for overexpression. (B-C) Control (*OK6-Gal4*) MN6/71b labelled with nc82 antibody (cyan, B) to mark brp positive active zones and merged with HRP (magenta, B’) which labels the axonal membrane. (C-C’) *dGgnbp2^OE-EPgy2^*(*OK6-Gal4/+;dGgnbp2^EPgy2^/ dGgnbp2^EPgy2^*) MN6/7 1b with the same labelling scheme as B. (D-G) Quantification of Control, *dGgnbp2^OE-Epgy2^* and *dGgnbp2^V5^*(*OK6-Gal4/+; UAS-dGgnbp2^V5^/ UAS-dGgnbp2^V5^*) bouton number normalised to muscle 6/7 surface area (1 x 10^3^μm^2^) (D) bouton size (E) brp puncta/bouton (F) and the number of brp puncta/neuromuscular junction (G). (H-I) Futsch labelling (green, H, H’) and merged with HRP (magenta, H’’, H’’’). High magnification of a terminal bouton showing a futsch loop (H’, H’’’). dGgnbp2^OE-EPgy2^ motor neuron with the same labelling scheme as in H (I-I’’’). (J-K) Quantification of Futsch in terminal boutons (J) and the different states of futsch labelling in control and *dGgnbp2^OE-EPgy2^* boutons (K). n=13-17. Quantification of compressed stacks done as in Figure 2. Each data point=average of 10 boutons. Data are represented as mean ± SEM. Depending on distribution of data an unpaired t test (D, J-K) or Mann-Whitney test (E-G). n.s *p*>0.05, \**p*<0.05, \*\*\**p*<0.001 and \*\*\*\**p*<0.0001. Scale bars denote 10um. See also Figure S4.

### Manipulating *dGgnbp2* levels disrupts locomotor performance in adult flies

The negative geotaxis assay is a common way to measure locomotor activity in *Drosophila*, taking advantage of the innate escape response when flies are tapped to the bottom of a tube and their ability to climb a set distance (8cm) in 10 seconds is quantified. Performance index is measured as the number of flies that cross this threshold over the total (Figure 4A). Human ALS-associated variants of *TDP-43* (*Tar DNA binding protein of 43kDa*) and *SOD1* (*Cu/Zn-superoxide dismutase 1*) expressed in *Drosophila* exhibit locomotor defects using this assay (*27*) (*28*).

**Figure 4.**
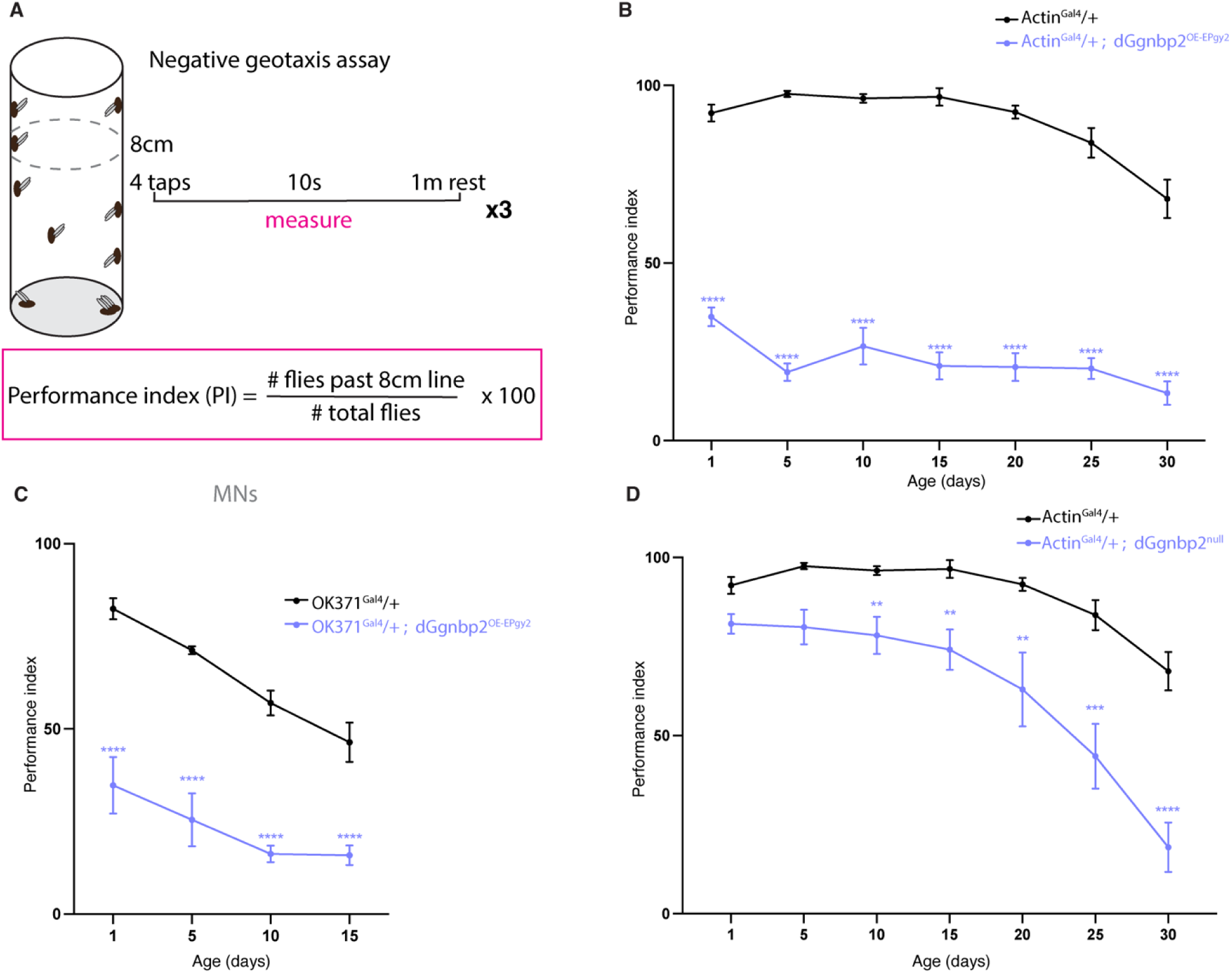
Manipulating *dGgnbp2* levels disrupts locomotor performance in adult flies. (A) Schematic of negative geotaxis assay. Flies were assessed 1-day post-eclosion and at 5-day intervals. (B) Control (*Actin-Gal4*, black) and *dGgnbp2^OE-EPgy2^* (*Actin-Gal4/+; dGgnbp2^OE-EPgy2^/ dGgnbp2^OE-EPgy2^,* purple) males were assessed for PI over 30 days. (C) Control (*OK371- Gal4*, grey) and *dGgnbp2^OE-EPgy2^*(purple) males assessed for PI over 15 days. (D) Control (*Actin-Gal4*, light grey) and *dGgnbp2^null^* (purple) assessed over 30 days. Flies were grouped into cohorts of 10 flies. Data are represented as mean ± SEM. Each datapoint represents the average of three technical replicates at each time point, per genotype. Three biological replicates (generated from three independent crosses) were carried out. n=7-9 cohorts of 10 flies. A Two-way ANOVA was performed with a Tukey’s or Sidak multiple comparison test. n.s *p*>0.05, \**p*<0.05, \*\*\**p*<0.001 and \*\*\*\**p*<0.0001. See also Figure S5.

When we drove *dGgnbp2* expression (*dGgnbp2^OE-EPgy2^*) in all cells using *Actin-Gal4*, a marked reduction in locomotor performance was evident over 30 days of testing. The phenotype in males (Figure 4B) was more severe than in females (Figure S5A), but both sexes exhibited defects even in young animals. To determine whether motor neurons were involved in this phenotype, we overexpressed *dGgnbp2* using a glutamatergic driver (*OK371-Gal4*) and observed a similar phenotype (Figure 4C, S5B). *dGgnbp2^null^*flies also showed a locomotor defect, but this was less severe and more age-dependent compared to *dGgnbp2^OE-EPgy2^* (Figure 4D, S5C). Due to previously reported locomotor defects in *w^1118^* flies (*29*) (Figure S5D-E), we used *Actin-Gal4* as our control and crossed this transgene into the *dGgnbp2^null^*background. Since both loss and gain of function for *dGgnbp2* lead to impairment, these results indicate that its expression needs to be tightly regulated for proper locomotion.

### *GGNBP2* is functionally conserved in flies

To test whether the human protein could rescue the fly phenotypes, we generated a *UAS* construct containing human *GGNBP2* with a C-terminal myc tag (*UAS-hGGNBP2^myc^*). We expressed this construct and a similar fly construct (*UAS-dGgnbp2^V5^*) (Figure 5A) with a motor neuron driver (*OK6-Gal4*) in a *dGgnbp2^null^* background and assessed motor neuron phenotypes. Remarkably, human *GGNBP2* was able to rescue all morphological defects including bouton number, bouton size and brp per bouton (Figure 5B-G, S6A). The human transgene driven by *Actin-Gal4* was also able to rescue locomotor defects observed in d*Ggnbp2^null^* animals (Figure 5H, S6B). Lastly, overexpression of the human transgene increased bouton size and the number of brp/bouton while reducing bouton number, similar to overexpression of *dGgnbp2* (Figure 5I- L, S6C-D). Together, these data indicate that GGNBP2 is highly conserved between humans and flies.

**Figure 5.**
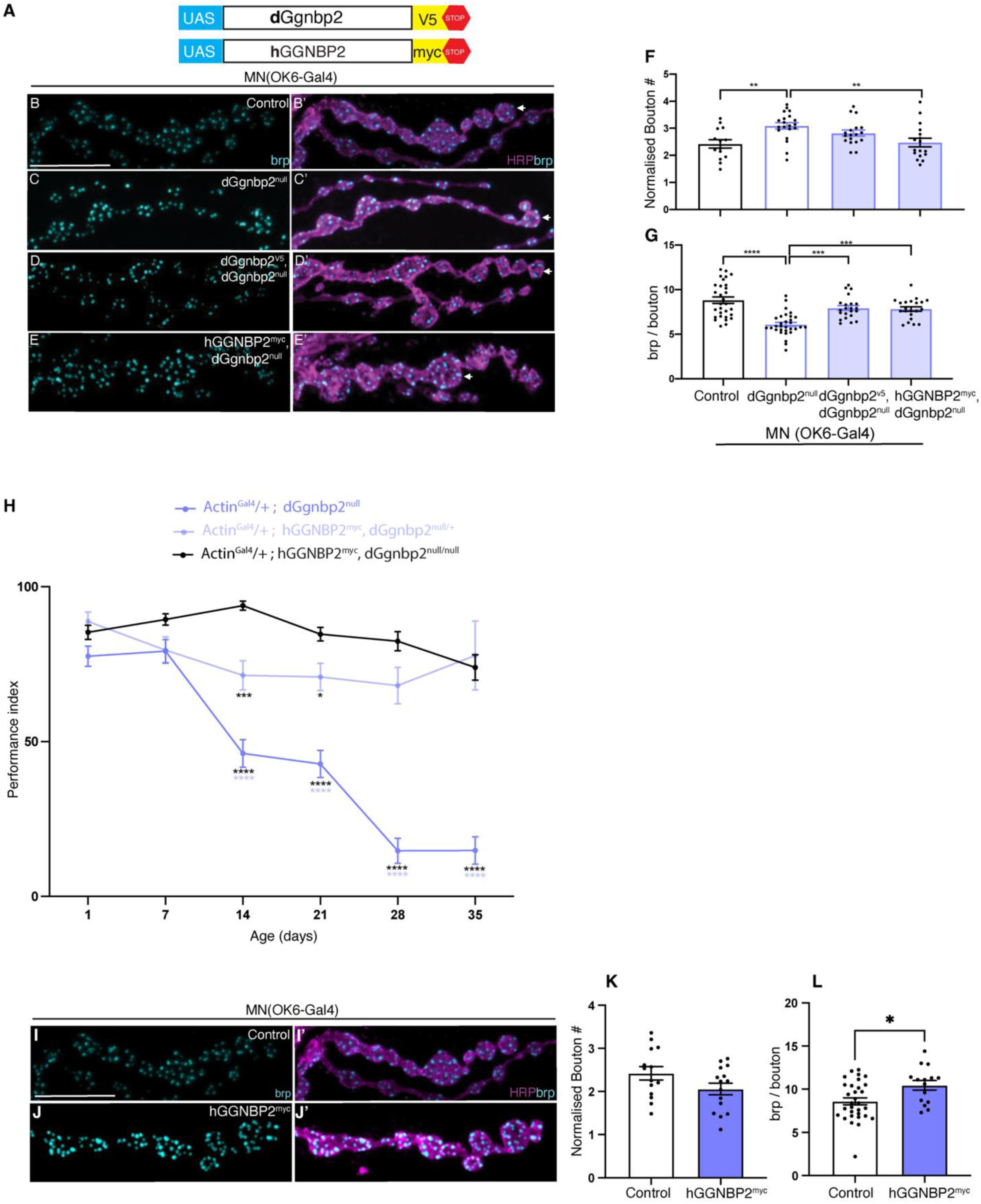
*GGNBP2* is functionally conserved in flies. (A) Schematic of *UAS-dGgnbp2^V5^*and *UAS-hGGNBP2^myc^*construct. (B-E) Rescue of *dGgnbp2^null^* motor neuron phenotypes with fly (D) and human (E) *GGNBP2*. Brp (cyan) and HRP (magenta) labeling of motor neurons as in previous figures. (F-G) Quantification of bouton number (F) and brp/bouton (G) in the four different genotypes: Control (*OK6-Gal4*), *dGgnbp2^null^, OK6-Gal4*; *dGgnbp2^V5^,* d*Ggnbp2^null^*/ d*Ggnbp2^null^, OK6-Gal4; hGGNBP2^myc^, dGgnbp2^null^*; *dGgnbp2^null^*. (H) Rescue of *dGgnbp2^null^*locomotion phenotype with *UAS hGGNBP2^myc^.* Males were assessed for PI over 35 days. (I-J) Overexpression of *hGGNBP2* in motor neurons with Brp (cyan) and HRP labeling. (K-L) Quantification of bouton number (K) and brp/bouton (L) in the genotypes: Control (*OK6-Gal4/+*) and *OK6-Gal4/+; hGGNBP2^myc^/+*. Quantification of compressed stacks as in Figure 2. Each data point=average of 10 bouton. Data are represented as mean ± SEM. An Ordinary one-way ANOVA with Tukey’s multiple comparison test (F-G), a Two-way ANOVA was performed with a Tukey’s or Sidak multiple comparison test (H) or unpaired t-test (K-L). n.s *p*>0.05, \**p*<0.05, \*\*\**p*<0.001 and \*\*\*\**p*<0.0001. Scale bars denote 10um. See also Figure S6.

### dGgnbp2 functions in the cytoplasm of motor neurons

Previous studies implicated Ggnbp2 in differentiation potentially as a transcriptional regulator (*4, 6*). To determine the localisation of fly dGgnbp2 protein, we knocked a V5-tagged version of the gene into the endogenous locus (*dGgnbp2^KI^*) using CRISPR homology-directed repair (HDR) (Figure 6A). The absence of morphologic and synaptic phenotypes at the NMJ of homozygous *dGgnbp2^KI^* animals demonstrated that the tagged protein is functional in motor neurons (Figure 6B-F, S7A). We visualized dGgnbp2 protein localization using a V5 antibody and identified motor neuron cell bodies in the ventral nerve cord with a *Gal4* driver expressed primarily in MN6/71b (*Exex-Gal4::CD8GFP*). The V5-tagged dGgnbp2 was not abundantly expressed anywhere in the larval ventral nerve cord or brain, and we could only detect it in larvae homozygous for the knock-in allele. In motor neurons, it was localized to the primarily to the cytoplasm (Figure 6G-H) but the possibility remains that it is present but undetectable in the nucleus.

**Figure 6.**
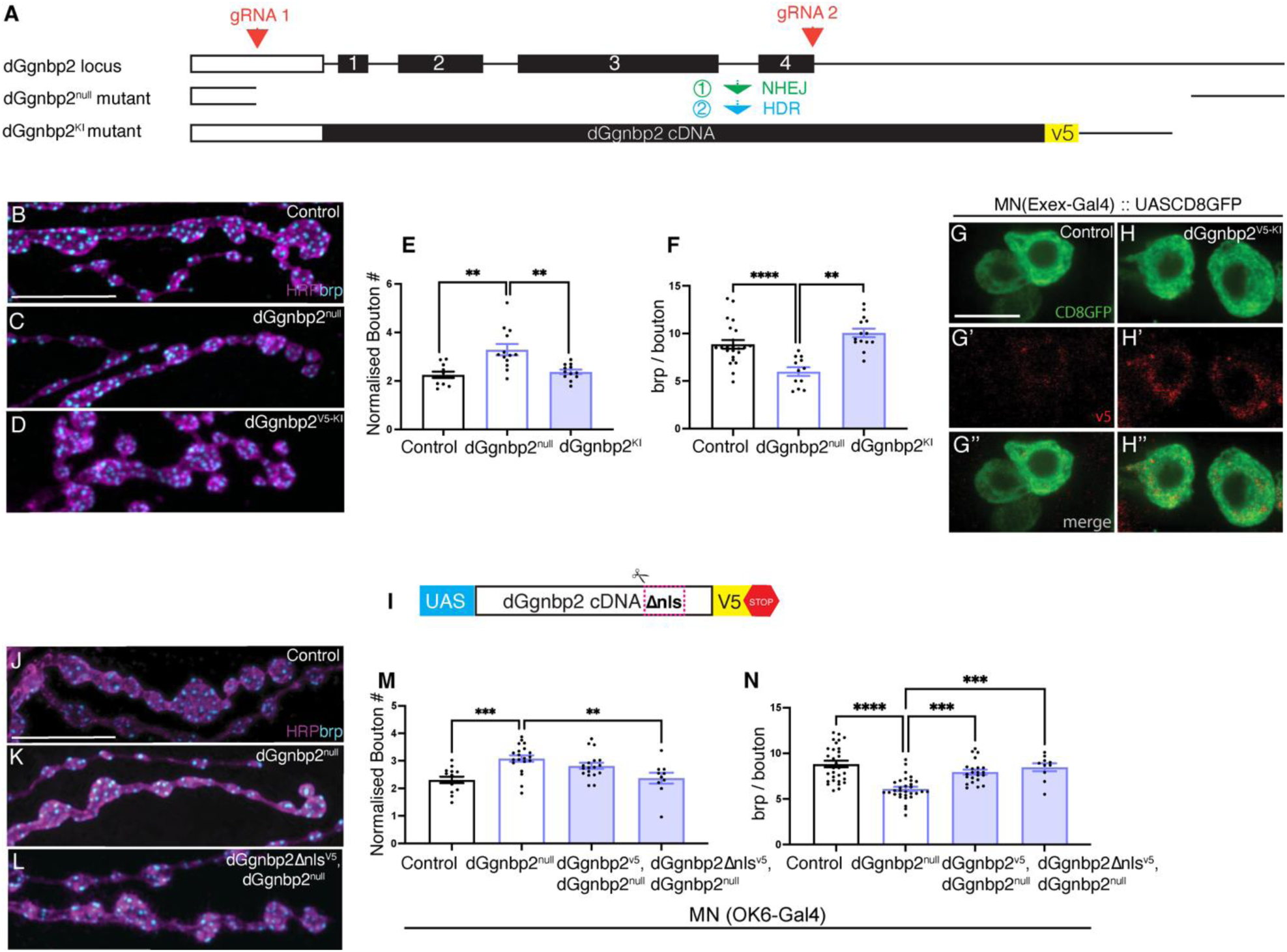
dGgnbp2 functions in the cytoplasm of motor neurons. Schematic of *dGgnbp2^V5-KI^* CRISPR knock-in, which was made using the *dGgnbp2^null^* strain. (B-D) Motor neurons labeled with brp and HRP as in previous figures. Control (*w^1118^*, B), *dGgnbp2^null^*(C), and homozygous *dGgnbp2^V5-KI^* (D). (E-F) Quantification of bouton and brp number as in Figure 2. (G-G’’) Control motor neuron cell bodies (*Exex-Gal4*::*UASCD8GFP)* labelled with GFP (G), V5 antibody (G’) and a merge (G’’). (H-H’’) Similar panel arrangement for flies expressing 2 copies of the *dGgnbp2^V5-KI^* construct. (I) Schematic of the UAS construct lacking the putative nuclear localization signal. (J-L) Merge of brp and HRP labeling in Control (*OK6-Gal4*, *J*), *dGgnbp2^null^* (K) and rescue animals (*OK6-Gal4/+; dGgnpb2*Δ*nls^V5^, dGgnbp2^null^,* L). (M-N) Quantification of bouton and brp number in animals rescued with WT and *dGgnpb2*Δ*nls^V5^.* n=9-25. Each data point=average of 10 boutons. Data are represented as mean ± SEM. Depending on distribution of data, an Ordinary one-way ANOVA with Tukey’s multiple comparison test (M-N) or a Kruskal-Wallis test with Dunn’s multiple comparison test was performed (G-H). n.s *p*>0.05, \**p*<0.05, \*\*\**p*<0.001 and \*\*\*\**p*<0.0001. Scale bars denote 10um. See also Figure S7.

To determine whether cytoplasmic dGgnbp2 is sufficient for its role in motor neurons, we identified a potential bipartite classical nuclear localization sequence (cNLS) using prediction software (*30-32*) and performed rescue experiments with a *UAS* transgene lacking this sequence (*UAS-dGgnpb2*Δ*nls^V5^*) (Figure 6I, S7B). This construct was able to completely rescue motor neuron phenotypes in *dGgnbp2^null^*animals (Figure 6J-N), confirming that these phenotypes are caused by a loss of cytoplasmic dGgnbp2.

### RNA-seq analysis of *Ggnbp2*

Given the proposed role for vertebrate *GGNBP2* in differentiation, we performed RNA-seq to investigate whether the protein is likely to regulate gene expression in the fly nervous system. RNA was collected from the third instar larval nervous system (brain and ventral nerve cord) from the following genotypes: *Control (w^1118^)*, *Actin-Gal4, dGgnbp2^null^*, *dGgnbp2^OE-EPgy2^*(*Actin::dGgnbp2^OE^*) and *dGgnbp2 ^EPgy2^ (insertion only)*, and three biological replicates were analysed for each. As anticipated, negligible levels of *dGgnbp2* were seen in *dGgnbp2^null^* and a∼7.5x increase in *dGgnbp2* was observed in *dGgnbp2^OE-EPgy2^* (Figure 7A-C). The *dGgnbp2^EPgy2^*insertion containing the *UAS* that we used for overexpression, appears to be a hypomorphic allele where a ∼50% decrease in *dGgnbp2* expression was observed. In *dGgnbp2^OE-EPgy2^* and *dGgnbp2^null^,* the number of differentially expressed genes and their respective fold changes were compared. Out of ∼5600 genes whose expression changed under the *dGgnbp2^OE-EPgy2^* and *dGgnbp2^null^* conditions, there were 83 (including *dGgnbp2*) with changes in opposite directions (Table S6), which would be predicted if *dGgnbp2* is regulating transcription. Several of the genes with the lowest p values are involved in biosynthetic processes such as fatty acid biosynthesis and serine transport into mitochondria. One gene, *gigas* (*gig*) encodes an mTOR inhibitor, which could be relevant to *dGgnbp2’s* role in PI3K signaling and autophagy (see below). However, the magnitudes of the changes in *gig* expression were small (10% increase with overexpression and a 10% decrease in the null allele); whether these are sufficient to drive biological effects or to implicate dGgnbp2 as a transcription factor is not clear. Overall, neither condition showed large changes in gene expression. In the portion of genes that were statistically significant for d*Ggnbp2^null^*and *dGgnbp2^OE-EPgy2^*, most exhibited modest fold changes of <1 in either direction (Figure S8A-F).

**Figure 7.**
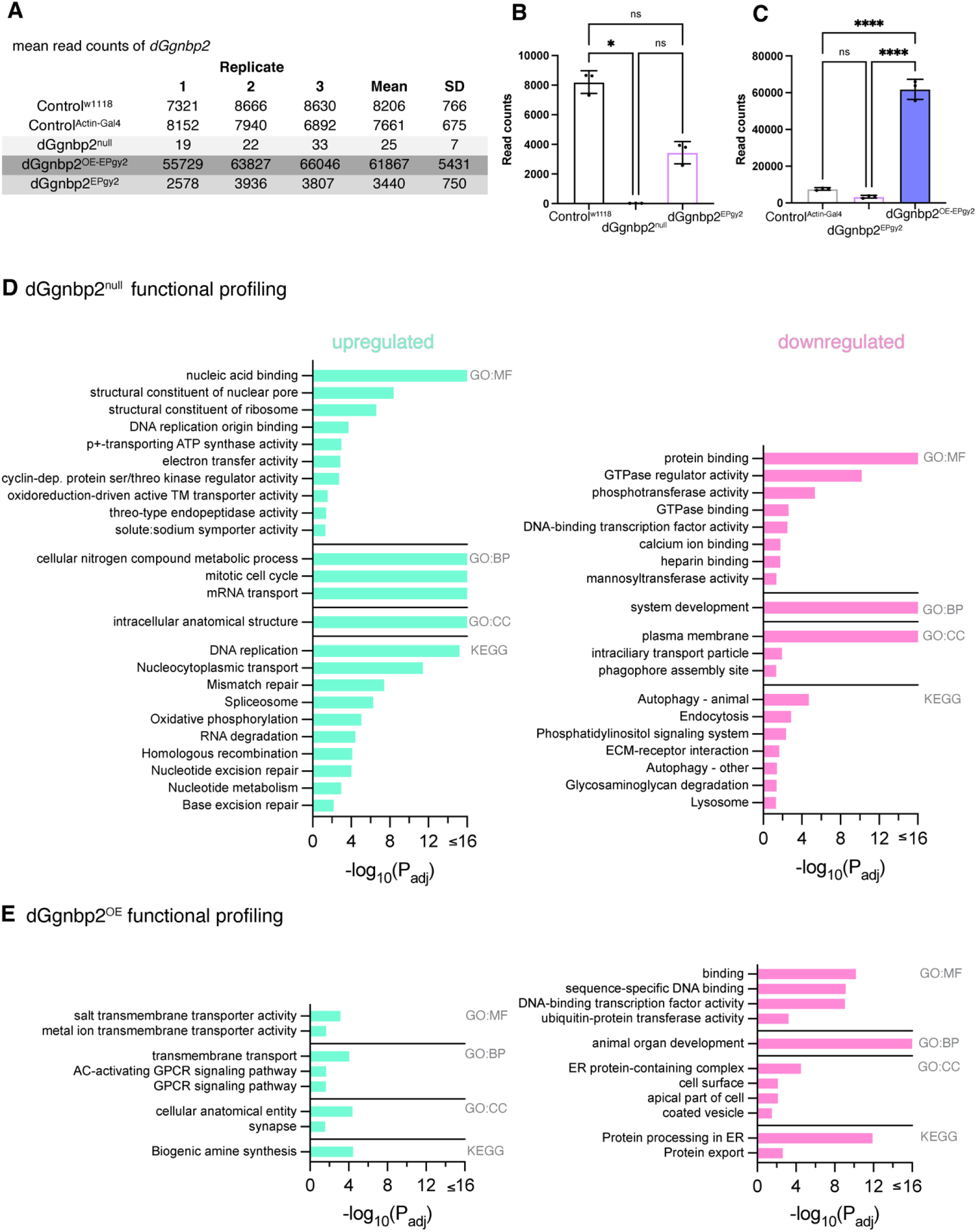
RNA-seq and functional profiling of *dGgnbp2*. (A) RNA-seq read counts of three replicates (including mean and SD) for *dGgnbp2* for two control conditions (*w^1118^* and *Actin-Gal4*) and three experimental *dGgnbp2^null^*, *dGgnbp2^OE-EPgy2^* and *dGgnbp2^EPgy2^* (hypomorphic). (B-C) Quantification of *dGgnbp2* read counts under lof (B) and gof conditions (C). (D-E) Functional profiling (gProfiler) for *dGgnbp2^null^*(D) and *dGgnbp2^OE-EPgy2^* (E) gene categories upregulated in green and downregulated in magenta. GO terms: molecular function (MF), biological process (BP), cellular component (CC), KEGG pathways (KEGG). Data are represented as mean ± SD. Depending on distribution of data an Ordinary one-way ANOVA with Tukey’s multiple comparison test (C) or Kruskal-Wallis test with Dunn’s multiple comparison test (B). n.s *p*>0.05, \**p*<0.05, \*\*\**p*<0.001 and \*\*\*\**p*<0.0001. See also Figure S8.

A functional enrichment analysis (g:GOSt) was performed using g:Profiler on differentially expressed genes that included GO (gene ontology) and KEGG pathway enrichment (*33*). In *dGgnbp2^null^*, there was an upregulation of nuclear factors and a down regulation of genes involved in autophagy, endocytosis and phosphatidylinositol signaling (Figure 7B). Although these data suggested that dGgnbp2 may have nuclear functions, the downregulation of cytoplasmic processes such as autophagy in the mutant suggested that it may also have functions outside the nucleus in neurons. In *dGgnbp2^OE-EPgy2^*there was an upregulation of transmembrane transporters and a downregulation of genes involved in DNA binding, organ development and ER processing (Figure 7C).

### *dGgnbp2* is linked to autophagy in motor neurons

Given the downregulation of autophagy, endocytosis and phosphatidylinosital signaling identified in *dGgnbp2^null^* animals by RNA-seq and its role in the cytoplasm of motor neurons we asked whether dGgnbp2 might contribute to autophagy, a cellular pathway that is commonly defective in ALS (Figure 8A). We used genetic interaction studies to determine whether *dGgnbp2* interacts with the fly orthologue of *TBK1*, called *ik2* in *Drosophila*. *TBK1* was originally identified as an ALS candidate gene by two independent studies through exome sequencing (*34, 35*). *TBK1* is a key regulator of autophagy and responsible for the phosphorylation of autophagy adaptors *p62*, *OPTN* (also ALS risk factors) and *NDP52* (*36*). Phosphorylation promotes complex formation with LC3-II and ubiquitinated cargo and thus enhances autophagy (*37*). *TBK1* has also been linked to another facet of autophagy where it is thought to be necessary for the maturation of autophagosomes to autophagolysosomes (*38*).Unlike the numerous gain of function/dominant mutations that have been identified in ALS, loss of function mutations in *TBK1* are prevalent (*35*). Since *lof* mutations are necessary for conducting genetic interaction studies, *TBK1* was an ideal candidate for testing a link with autophagy.

**Figure 8.**
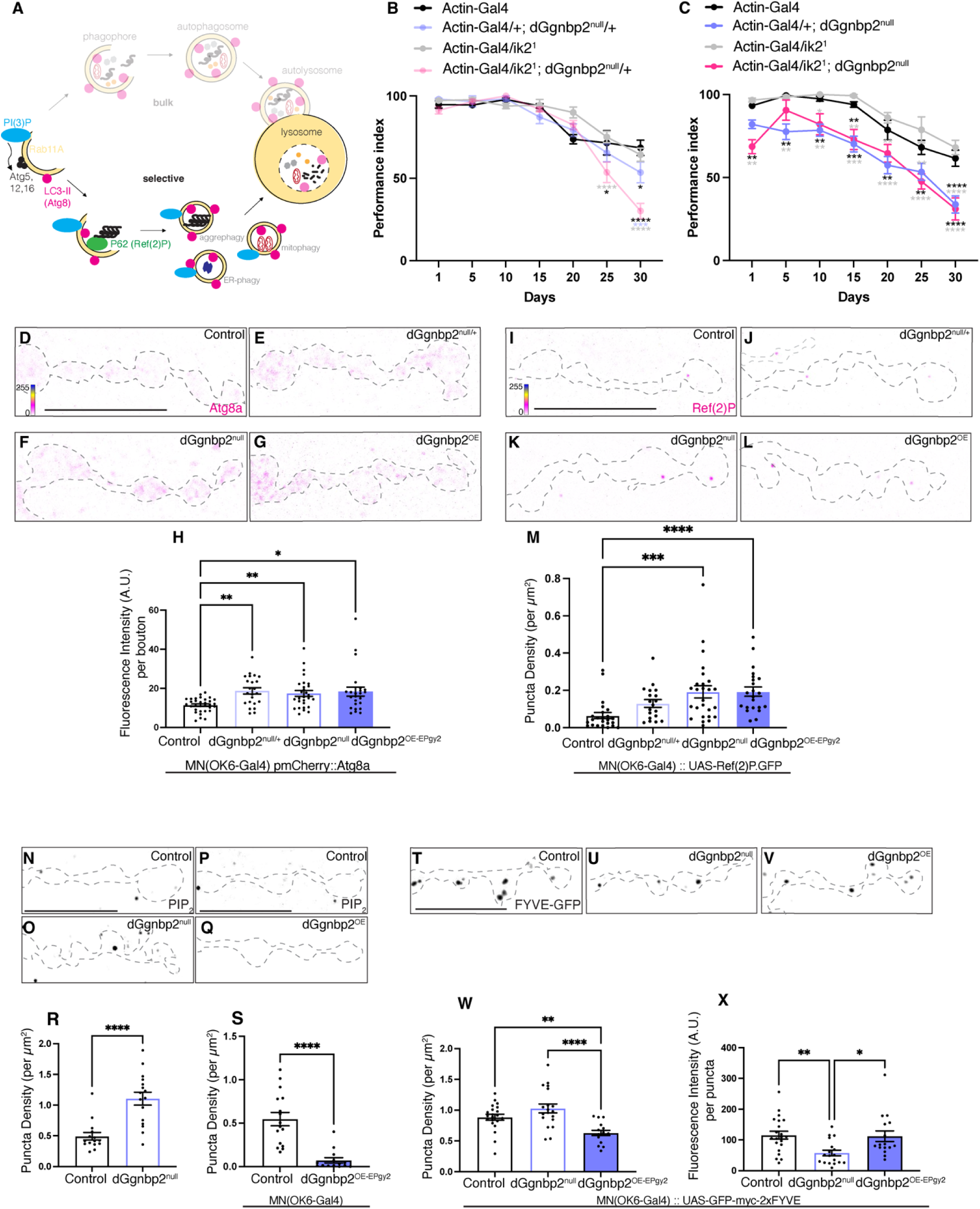
*dGgnbp2* levels are critical for autophagy. (A) Schematic of bulk and selective autophagy. (B-C) Genetic interactions between *dGgnbp2* and *ik2* assessed through negative geotaxis as in Figure 4. Heterozygous combinations (B) and *ik2*/+; *dGgnbp2^null^* combinations (C) of males were assessed over 30 days. n=9-11 cohorts of 10 flies. A Two-way ANOVA was performed with a Tukey’s or Sidak multiple comparison test. n.s *p*>0.05, \**p*<0.05, \*\*\**p*<0.001 and \*\*\*\**p*<0.0001. (D-G) MN6/71b were labelled with HRP (not shown) and motor neuron terminal outline is overlayed (grey, dotted) on images expressing endogenous *pmCherry-Atg8a* (4 LUT magenta – blue) in different *dGgnbp2* genotypes: Control *dGgnbp2^+/+^* (D), *dGgnbp2^null/+^*(E), *dGgnbp2^null^* (F), *dGgnbp2^OE-EPgy2^* (G). All genotypes were in a *OK6-Gal4* background. (H) Quantification of fluorescence intensity (A.U.) per bouton for Atg8a. n=21-24. (I-L) MN6/71b labelled with HRP (not shown) and motor neuron terminal outline is overlayed (grey, dotted) on images expressing *OK6::Ref(2)P.GFP* in different *dGgnbp2* genotypes: Control *dGgnbp2^+/+^* (I), *dGgnbp2^null/+^*(J), *dGgnbp2^null^* (K), *dGgnbp2^OE-EPgy2^* (L). (M) Quantification of Ref(2)P puncta density (per μm^2^). n=19-21. (N-Q) MN6/71b labelled with HRP (not shown) and motor neuron terminal outline is overlayed (grey, dotted) on images for an antibody against PIP_2_ (black puncta) in Control (*w^1118^*, N), *dGgnbp2^null^* (O), Control (*OK6-Gal4*, P), *dGgnbp2^OE-EPgy2^* (Q). (R-S) Quantification of PIP_2_ puncta density (per μm^2^). n=13-16. (T-V) MN6/71b labelled with HRP (not shown) and motor neuron terminal is overlayed (grey, dotted) on images expressing the PI(3)P reporter *OK6::GFP-myc-2xFYVE* (black puncta) in the following *dGgnbp2* genotypes: Control *dGgnbp2^+/+^*(T), *dGgnbp2^null^* (U) and *dGgnbp2^OE-EPgy2^* (V). (W-X) Quantification of *GFP-myc-2xFYVE* puncta density (per μm^2^) and *GFP-myc-2xFYVE* puncta fluorescence intensity (A.U.). n=12-19. Each data point=average of 10 boutons. Data are represented as mean ± SEM. Depending on distribution of data a Mann-Whitney test (R-S) or an Ordinary one-way ANOVA with Tukey’s multiple comparison test (W-X) or Kruskal-Wallis test with Dunn’s multiple comparison test (H, M). n.s *p*>0.05, \**p*<0.05, \*\*\**p*<0.001 and \*\*\*\**p*<0.0001. Scale bars denote 10um. See also Figure S9.

We obtained a *lof* allele for *ik2* (*ik2^1^*) that is homozygous lethal early in larval development. To assess genetic interactions between *ik2* and *dGgnbp2*, we combined the two heterozygous mutations and asked whether they displayed *dGgnbp2^null^*phenotypes. Heterozygotes for either *ik2* or *dGgnbp2* alone showed no observable morphological or synaptic defects in motor neurons. In contrast, double-heterozygotes displayed motor neuron defects where both bouton size and brp number was greatly reduced (Figure S9A-C). This suggested that these genes could function in the same genetic pathway. To confirm this, we also assessed locomotion performance of double heterozygotes using negative geotaxis. In this assay, *dGgnbp2* heterozygotes (*dGgnbp2^null/+^*) showed a slight defect at Day 30 whereas *ik2^1/+^*animals performed similar to controls. Double heterozygotes (*ik2^1/+^; dGgnbp2^null/+^*) initially behaved similar to controls but exhibited a significant decline at the 25- and 30-day time points, more than either heterozygote alone (Figure 8B), suggesting a common pathway. If *dGgnbp2* and *ik2* function in the same linear pathway, then a homozygous null mutation in *dGgnbp2* should disable this pathway and removing one allele of *ik2* should not modify the phenotype. This is indeed what we observed; in a *dGgnbp2^null^* background, *ik2^1/+^* exhibited a similar locomotion defect to *dGgnbp2^null^* alone (Figure 8C). However, the same was not true in motor neurons as *ik2^1/+^* suppressed the *dGgnbp2^null^* phenotypes (Figure S9A-C), suggesting a more complicated relationship between *dGgnbp2* and *ik2* (*TBK1*) in regulating motor neuron development.

### *dGgnbp2* levels are critical for autophagy

Autophagy is a process that maintains cellular homeostasis by removing damaged organelles and macromolecules through fusion with lysosomes (*39*). To directly test whether *dGgnbp2* is involved in this process, autophagy markers Atg8 (LC3 orthologue) and Ref(2)P (p62 orthologue) were assessed in motor neurons of different *dGgnbp2* genotypes. Atg8 is localised at the autophagosome membrane and is important for cargo recruitment and biogenesis (*40, 41*). We used an endogenously tagged pmCherry-Atg8a to assess Atg8a levels using fluorescence intensity. To make the genetics consistent, we worked in an *OK6-Gal4* background for all *dGgnbp2* genotypes, even if the *Gal4* was not driving a *UAS* construct. In control motor neuron boutons, endogenous pmCherry-Atg8a was diffuse and weakly expressed. In contrast, the manipulation of *dGgnbp2* levels (*dGgnbp2^null/+^*, *dGgnbp2^null^*and *dGgnbp2^OE-EPgy2^)* led to an increase in Atg8a fluorescence (Figure 8D-H). The accumulation of Atg8a could mean either that autophagy induction is increased or that the process is blocked/delayed downstream of autophagosome formation (*42*).To distinguish between these possibilities, we overexpressed GFP-tagged Ref(2)P in motor neurons and asked whether this protein also accumulates. Like its orthologue p62, Ref(2)P delivers cargo to autophagosomes and itself is degraded in the process (*43-46*). Ref(2)P formed sparse puncta in control motor neurons (Figure 8I). When *dGgnbp2* was decreased or increased, Ref(2)P puncta density increased, although it was not statistically significant in the heterozygotes (Figure 8J-M, S9D). These data suggest that both increases and decreases in *dGgnbp2* expression inhibit autophagy.

### *dGgnbp2* alters autophagy-relevant phospholipids

Human *GGNBP2* has been implicated as a tumor suppressor in both brain and breast cancer, although the proposed mechanisms are distinct (Zhan 2017; Lan 2016). One of the growth promoting pathways that GGNBP2 has been proposed to inhibit is the PI3K/Akt pathway (Lan 2016). PI3K plays a crucial role in autophagy so we investigated whether lipid phosphorylation is modified in motor neurons lacking or overexpressing *dGgnbp2*.

Autophagy has a complicated relationship with PI3K signaling. The PI3K/Akt/mTor pathway, driven primarily by class I PI3K on the plasma membrane, inhibits autophagy through mTOR (*47*). In contrast, the class III PI3K that generates PI(3)P (Vps34) is an integral component of the autophagy machinery. PI(3)P present on the autophagosome membrane is necessary for recruiting autophagy effectors (*47-49*).

To test whether *dGgnbp2* levels affect PI3K signaling in motor neurons, we used antibodies and a fluorescent reporter to measure two species of phosphorylated lipid. An antibody to PI(4,5)P2 (PIP_2_) was used as a readout for class I PI3K activity (*50*). Increased levels of this substrate indicate a reduction in enzyme activity, whereas a reduction in PIP_2_ reflects increased class I activity. To ensure that the PIP_2_ antibody could detect these readouts, we assessed PIP_2_ puncta in motor neurons that overexpressed class I PI3K (PI3K^92E^) and a ‘kinase-dead’ PI3K (PI3K^KD^) (Figure S9E-G). PIP2 puncta density was quantified and found to be decreased upon PI3K overexpression as expected (Figure S9H).

In control motor neurons, endogenous PIP_2_ was punctate so we measured the number of puncta per μm^2^. Surprisingly, PIP_2_ puncta density was increased in *dGgnbp2^null^*compared to control (Figure 8N-O, R, S9I), suggesting a reduction in class I PI3K signaling. Consistent with this, overexpression of *dGgnbp2* decreased PIP_2_ puncta density, suggesting that it promotes PI3K signaling (Figure 8P-Q, S, S9J). This contrasts with the studies in cancer cell lines that implicated GGNBP2 as a tumor suppressor that inhibits the PI3K/Akt pathway. Instead, our data are consistent with dGgnbp2 promoting class I PI3K in motor neurons.

To quantify lipid species derived from class III PI3K, we expressed the fluorescent reporter that binds directly to PI(3)P, 2xFYVE-GFP, in motor neurons (*51*). There should be a direct relationship between FYVE-GFP puncta and PI(3)P levels, which indirectly assesses the activity of class III PI3K. The PI(3)P reporter (*UAS-GFP-myc-2xFYVE*) generated large puncta in control neurons. The density of PI(3)P puncta was unchanged in *dGgnbp2^null^* animals, but was reduced when *dGgnbp2* was overexpressed (Figure 8T-W), suggesting that excessive dGgnbp2 inhibits class III PI3K. Although the number of puncta was unchanged in *dGgnbp2^null^*animals, the intensity of individual puncta appeared reduced. We therefore quantified the average puncta fluorescent intensity in each genotype and found that it was indeed reduced in *dGgnbp2^null^* motor neurons (Figure 8X). These reductions in PI(3)P in *dGgnbp2^null^* and *dGgnbp2^OE-EPgy2^*are consistent with changes in the level of this protein negatively affecting autophagic flux, potentially by interfering with class III PI3K activity or by stimulating myotubularin phosphatases that convert PI(3)P to PI.

## Discussion

Our investigation in *Drosophila* prioritizes further studies into *GGNBP2* as a risk gene for ALS within the associated genomic locus. The SNP associated with ALS on chromosome 17 implicates 5 closely linked genes (*GGNBP2*, *PIGW*, *MYO19, DHRS11* and *ZNHIT3*) but both a gene-based test and changes in gene expression associated with this risk allele implicate *GGNBP2* as the top candidate in this region (Figure 1). Our functional data in flies support this conclusion. We demonstrate that both loss and gain of function *dGgnbp2* mutants exhibit morphological and synaptic phenotypes in motor neurons. We show through rescue experiments that the human and fly *GGNBP2* genes are functionally conserved in motor neurons. Overexpression of *dGgnbp2* causes a severe climbing defect in adult flies, linking the function of this gene to motor output. In addition, we provide the first mechanistic clues about how this protein functions in motor neurons; it regulates autophagy, a process that is frequently perturbed in ALS (*52*). *dGgnbp2* genetically interacts with *ik2* (human *TBK1*), a kinase that phosphorylates proteins involved in autophagosome formation, and impacts PI3K-related pathways that regulate autophagy.

We demonstrate in flies that overexpression of *dGgnbp2* severely impacts fly locomotion and motor neuron autophagy, two phenotypes associated with ALS. Loss of function data for fly *dGgnbp2* are also consistent with human data that have implicated this gene in other neurological disorders and suggest that it is limiting in the nervous system. In humans, there is selection against null mutations in *GGNBP2* where the observed/expected ratio of homozygous loss of function mutations is 0/40.8, n=141,456, pLI=1 (gnomAD v2.1.1) (*53*). Interestingly, rare heterozygous loss of function variants in *GGNBP2* have been reported in two autism cohorts (*54*) (*55*) and thus could indicate a role for *GGNBP2* in brain development. Additionally, this chromosomal region (1.4Mb, 17q12, spanning 15 genes) is a known risk region for both autism and schizophrenia, with the deletion identified in 18 patients and no controls (n=15,749) (*56*). Although fly *dGgnbp2* is recessive for developmental phenotypes in motor neurons (Figure S3), heterozygotes exhibit autophagic flux phenotypes (Figure 8). Thus, reductions and increases in the expression of this gene lead to phenotypes in both humans and flies.

One difference between overexpression of *dGgnbp2* in flies and humans with progressive neurological diseases such as ALS is that the locomotion phenotype in flies is severe in young adults (Figure 4), suggesting that the phenotype may be predominantly developmental. Even in females where the phenotype was less severe (Figure S5), there were clear deficits at the early time points. Our phenotypes in larval motor neurons overexpressing *dGgnbp2* are consistent with this. Significant changes in motor neuron morphology and synaptic makeup demonstrate that excessive *dGgnbp2* (7.5x, Figure 7) impairs the development of these neurons. It is possible that a more modest increase in *dGgnbp2* may not affect development and lead to an age-dependent decline in locomotor performance.

In contrast to its role in reproductive tissues such as sperm and trophoblast cells where it regulates nuclear processes (*4, 6*), our data demonstrate that dGgnbp2 functions in the cytoplasm of motor neurons (Figure 5). Sperm contain a germ cell specific protein called gametogenetin (GGN) that binds to GGNBP2 (*3*), and the complex has been implicated in DNA repair (*5*). In trophoblast stem cells of the placenta, GGNBP2 is in the nucleus, necessary for differentiation, and implicated in transcriptional repression (*6*). Consistent with our data arguing for a cytoplasmic role for GGNBP2, a recent study into the CCR4-NOT1 mRNA deadenylase complex, which also functions in the cytoplasm, identified GGNBP2 as a protein binding partner (*19*). These studies were carried out in cell lines, so it remains to be determined whether GGNBP2 contributes to this complex in neurons. Our RNA-seq data identified 36 transcripts whose expression increased with *dGgnbp2^OE-EPgy2^* and decreased in the *dGgnbp2^null^*and 47 transcripts whose levels showed the opposite pattern (Table S6). This indicates that dGgnbp2 has the potential to regulate mRNA stability through the CCR4-NOT complex in neurons. However, there were many more differentially expressed genes that did not show inverse patterns with the two genetic manipulations and as outlined above for *gig*, most of the changes in expression were small.

An analysis of differentially expressed genes from RNA-seq data that included GO and KEGG pathway enrichment revealed a downregulation of autophagy, endocytosis and PI signaling in *dGgnbp2^null^* animals (Figure 7). Consistent with *dGgnbp2* regulating autophagy, it genetically interacted with *ik2* (*TBK1*), a kinase that regulates autophagosome formation (*36*). Double heterozygotes exhibited synergistic phenotypes in motor neuron development (Figure S9) and climbing performance (Figure 8) compared to either heterozygote alone. This suggested that these two genes act in the same pathway. However, in the absence of *dGgnbp2*, the *ik2* heterozygotes suppressed motor neuron phenotypes, rather than leaving them unchanged, as was observed for locomotor performance. This argues that the relationship between these genes during development is likely more complicated than a simple linear pathway. We used two markers for autophagy to demonstrate that this process is blocked/delayed in animals lacking or overexpressing *dGgnbp2*. Both Atg8a (LC3) and Ref(2)P (p62) accumulated when we removed 1 or 2 copies of *dGgnbp2* and when we overexpressed it. This suggests that autophagic flux is impaired in response to changes in dGgnbp2 levels. We also demonstrated that *dGgnbp2* regulates phosphorylated species of phosphatidylinositol in different genotypes. Our results contrast those from cancer studies demonstrating that GGNBP2 suppresses class I PI3K in glioma cell lines (*57*). We observed an increase in PIP_2_ in *dGgnbp2^null^* animals and a decrease in PIP_2_ when *dGgnbp2* was overexpressed (Figure 8). These data indicate that in motor neurons, *dGgnbp2* promotes conversion from PIP_2_ to PIP_3_, potentially by regulating class I PI3K activity, which would inhibit autophagy induction. Analysis of PI(3)P levels in different genotypes also exhibited a *dGgnbp2* dose-dependent effect, but in this case both a reduction and an increase in *dGgnbp2* led to a decrease in PI(3)P (Figure 8).

For larval motor neuron morphology phenotypes, loss and gain of function of *dGgnbp2* showed inverse phenotypes. For autophagy, however, both conditions impaired this cellular process rather than one increasing it and the other decreasing it. Although we are unsure why this is the case, it is not unprecedented. For example, both loss and gain of function of *Caz*, the fly homologue of the ALS gene *FUS*, exhibit similar phenotypes (*58*). Similarly, dopamine dosage follows an inverted U-shaped curve in terms of cognitive performance with both too little and too much showing poor performance (*59*). Although it is speculative, we propose that *dGgnbp2* is a limiting factor in autophagy that is part of a larger protein complex. For example, *dGgnbp2* could be involved with the assembly of PI(3)P-positive autophagosomes, where it might interact with the class III enzyme (Vps34) present on these membranes. Reductions in the protein would therefore impair the function of this complex and too much *dGgnbp2* could do the same through gain of function mechanisms that sequester other limiting components and therefore impair complex function.

This study investigated the gene most likely to contribute to ALS risk on chromosome 17 using *Drosophila*. We have provided strong evidence for a role for *dGgnbp2* in motor neurons and pathways linked to ALS pathology. However, one limitation of this study is that other genes in this region could also be contributing to ALS risk. Clarification of this point will arise from future GWAS and expression studies that encompass larger ALS cohorts. We note a recent study that analyzed spinal cord expression data in ALS cases and controls found that *GGNBP2* was significantly upregulated in the ALS cohort and that *GGNBP2* expression (but not other linked genes in the locus) negatively correlated with duration of disease (*60*). While we demonstrated conservation between fly and human *GGNBP2*, further investigation into how altering the levels of this gene affect human motor neuron development and function is needed.

In conclusion, we have demonstrated that *dGgnbp2* is required in motor neurons, functionally conserved between flies and humans and a regulator of autophagy. Changes in lipid phosphorylation represent a potential mechanism for the impairment of autophagy. Overexpression of *dGgnbp2* decreases PIP_2_ levels, consistent with an increase in class I PI3K activity that would activate mTOR and inhibit autophagy. This same condition also decreases the number of PI(3)P-positive autophagosomes, thereby inhibiting autophagy at a later step and through a distinct mechanism. Given that autophagy is commonly impaired in ALS, reducing GGNBP2 levels could be one strategy for promoting this crucial cellular process for therapeutic benefit.

## Materials and Methods

### GWAS and post-analyses of human data

Using the largest ALS GWAS (*12*) (N_cases_=29,612 N_controls_=122,656, multi-ancestry) we conducted analyses integrating functional and biological information to support gene prioritisation from the Chromosome 17 locus. Given the high-LD in the region we first used the GCTA-COJO method (*13*). This can determine if the most associated SNP accounted for the association signal in the region (other SNP associations reflecting correlation to the most associated SNP), or if conditioning on the most associated SNP would identify secondary significantly associated SNPs using a COJO *p* value threshold of *p*<5×10^-7^.

We then used gene-based tests including mBAT-combo (*14*). This is a recently developed test better powered than other methods (i.e. fastBAT) to detect multi-SNP associations as it accounts for masking effects – common in complex traits/disease. We applied this to the ALS GWAS summary statistics. Only the European samples were used (N_cases_=27,205 N_controls_=110,881)(*12*) as the correlation structure between SNPs is determined from a reference sample of matched ancestry. Input SNPs were mapped to 18,689 protein-coding genes with a 0kb window and genome-wide significance defined at P = 0.05/18689 = 2.68e-6.

We next ran a summary-based Mendelian randomization (SMR) analysis (*61*). This approach can test if the effect of a SNP on the phenotype is mediated by gene expression and therefore, prioritize genes underlying GWAS for functional studies. We used expression eQTLs (SNPs associated with gene expression) data from two large meta-analyses. The blood expression data (eQTLGen, n = 31,648, n_eQTL_=16,987, (*62*)) included data from 37 cohorts (mostly European ancestry) not enriched for any disease. The brain expression data (MetaBrain, n = 2,970, n_eQTL_=12,307 (*63*)) was from the cortex of 14 datasets also not enriched for any disease. Given that the *GGNBP2* is a common allele, we expect ∼25% of individuals to be homozygous for this SNP. The Bonferroni corrected significance thresholds were p=0.05/ n_eQTL_ for blood (discovery) and p=0.05/5 for brain (replication).

Given rare and common variants have been identified to converge on the same gene (*12*), rare-variant burden testing (MAF <1 and 0.5%, disruptive + damaging variants) was examined using Project MinE whole genome sequences (N_cases_=4,633 N_controls_=1,832) (*64*). Firth logistic regression was used to compare cases vs. controls.

### Fly genetics and transgenic lines

Flies were cultured on standard yeast-agar media at 25°C in a room with windows that exposed them to natural day/night cycles. The following *Gal4* driver lines, transgenes for genetic analyses and transgenic generation were used:

| Reagent<br>Transgenic lines | Source | Identifier |
| --- | --- | --- |
| *OK6-Gal4* | Bloomington | BL#64199 |
| *BG487-Gal4* | Bloomington | BL#51634 |
| *elav-Gal4* | Bloomington | BL#8765 |
| *Actin-Gal4* | Bloomington | BL#25374 |
| *OK371-Gal4* | Bloomington | BL#26160 |
| *CG2182<sup>EPgy2</sup>* | Bloomington | BL#15311 |
| *ik2<sup>1</sup>* | Bloomington | BL#5322 |
| *Df(3R)BSC464* | Bloomington | BL#24968 |
| *Df(3R)BSC47* | Bloomington | NA |
| *Actin-Cas9* | Bloomington | BL#54590 |
| *Actin-Cas9, Lig4* | Bloomington | BL#58492 |
| *M{3xP3-RFP.attP}ZH-86Fb* | Bloomington | BL#24749 |
| *pmCherry::Atg8a* | Gift from Donna Denton,<br>University of South Australia | NA |
| *UAS-GFP.Ref(2)P* | Thomas Neufeld (65) | NA |
| *UAS-GFP-myc-2xFYVE* | Bloomington | BL#42712 |
| *UAS-PI3K<sup>92E</sup>* | Bloomington | BL#8286 |
| *UAS-PI3K<sup>KD</sup>* | Bloomington | BL#8289 |

### Molecular Biology Cloning

The fly and human *GGNBP2* cDNAs were synthesized by General Biosystems in a pBluScriptII SK+ vector and PCR was used to generate tagged versions of the two genes, which were then cloned into the pMUH-JFRC plasmid (Addgene) using Gibson cloning (NEB). Similarly, the Dnls construct was generated using PCR and Gibson cloning. Listed below: *UAS- dGgnbp2^V5^*, *UAS-hGGNBP2^myc^* and *UAS-dGgnbp2*Δ*nls^V5^*.

### CRISPR gene editing

*dGgnbp2^null^* allele was generated using two guide RNAs (gRNAs) to completely remove the coding region of *dGgnbp2* via NHEJ. This was confirmed through sanger sequencing.

Embryos (*Actin-Cas9*) were co-injected with Ebony to assist with screening (*66*).

gRNA^1^ *dGgnbp2^null^*– 5’ GTTCTATGTCAAACACGTAGT 3’
gRNA^2^ *dGgnbp2^null^*– 5’ GAAGCGGACGACTTCTGCAC 3’

A d*Ggnbp2* knockin (*dGgnbp2^KI-V5^*) allele was generated via CRISPR HDR from the *dGgnbp2^null^* fly line using the guide below. A V5-tagged version of fly *dGgnbp2* was generated by PCR from the same pBluScript CG2182 vector described above which contained 5’ and 3’ homologous arms to facilitate HDR. Template and gRNAs were co-injected into embryos (*Actin-Cas9, Lig4; Ggnbp2^null^*) and confirmed through sanger sequencing.

NC062 – 5’ GTTTCATTGGCAAGTCGGTC 3’

All gRNAs were designed using CRISPR Optimal Target Finder (*67*).

### Immunohistochemistry

Flies were maintained on cornmeal/agar media at 25°C and were dissected for analysis within one day of eclosion. Third instar larvae were filleted/dissected in ice cold PBS and fixed in 4% PFA for 20 minutes. Larval preparations for NMJ analysis were blocked in PBS containing 10% goat serum, 0.1% Triton X-100 for 60 minutes within an hour of fixation. Larval preparations for VNC analysis were blocked in PBS containing 10% goat serum, 0.5% Trition X-100 for 60 minutes.

Primary and secondary antibodies were incubated overnight and washed 3X with PBS. Antibody dilutions used were as follows: mouse anti-nc82 (1:100, DSHB), mouse anti-22c10 (1:100, DSHB), mouse anti-GluRIIA (8B4D2 (MH2B)) (1:100, DSHB), mouse anti-LC28 (1:100, DSHB), chicken anti-GFP (1:1000, Abcam), mouse anti-PIP_2_ (1:500, Abcam), DyLight anti-mouse 647(1:1000, Jackson Laboratory), Alexa Fluor 647 anti-mouse IgM mu chain (1:2000, Abcam), Alexa Flour 488 anti-chicken (1:1000, Abcam), anti-HRP-conjugated (Cy3 or Alexa Fluor 647, Jackson Laboratory), V5-tag:DyLight anti-mouse 550 (1:500 AbD Serotec). Hoescht (1:20 000, 33258 abcam). Samples were analysed within a week of IHC.

| Reagent Antibodies | Source | Identifier |
| --- | --- | --- |
| mouse anti-nc82 | DSHB | AB_2314866 |
| mouse anti-22c10 | DSHB | AB_528403 |
| mouse anti-GluRIIA (8B4D2 (MH2B)) | DSHB | AB_528269 |
| mouse anti-LC28 | DSHB | AB_528339 |
| chicken anti-GFP | Abcam | (ab13970) |
| mouse anti-PIP<sub>2</sub> | Abcam | (ab11039) |
| DyLight anti-mouse 647 | Jackson laboratory | AB_2338907 |
| Alexa Fluor 647 anti-mouse IgM mu chain | Abcam | (ab150123) |
| Alexa Flour 488 anti-chicken | Abcam | (ab150169) |
| anti-HRP-conjugated-Cy3 | Jackson laboratory | AB_2338959 |
| anti-HRP-conjugated-Alexa Fluor 647 | Jackson laboratory | AB_2338967 |
| V5-tag:DyLight anti-mouse 550 | BioRad | MCA1360D550GA |
| Hoescht | Abcam | (ab228551) |

### Image acquisition and deconvolution

Imaging was performed at the Queensland Brain Institute’s Advanced Microscopy Facility and School of Biomedical Sciences Imaging Facility. Data was imaged on a spinning-disk confocal system (Marianas; 3I, Inc.) consisting of a Axio Observer Z1 (Carl Zeiss) equipped with a CSU-W1 spinning-disk head (Yokogawa Corporation of America), ORCA-Flash4.0 v2 sCMOS camera (Hamamatsu Photonics), 63x 1.4 NA P-Apo, 100x 1.3 NA P-Apo and 150x 1.3 NA P-Apo objectives. Image acquisition was performed using SlideBook 6.0 (3I, Inc). Ggnbp2 localisation and GluR was visualised and imaged using a Nikon Plan Apo Lambda 100x/1.45 NA oil-immersion objective on a spinning disk confocal microscope (Diskovery; Andor Technology, UK) built around a Nikon Ti-E body (Nikon Corporation, Japan) and equipped with two Zyla 4.2 sCMOS cameras (Andor Technology) and controlled by Nikon NIS software.

### Image Analysis

### Quantification of synaptic morphology at the third instar larval NMJ

HRP labelling was used as a guide to delineate boutons at the NMJ. Analysis was performed using ImageJ. Three measurements were used (1) the number of brp puncta per bouton in 1b, (2) the size (surface area in μm^2^) of 1b and (3) the number of combined (1b and 1s) boutons normalised to muscle surface area (surface area in μm^2^). Ten boutons were analysed for (1) and (2) and averaged. Imaris (Bitplane, St. Paul, MN) was used to quantify synapse number per NMJ (shown as puncta) after manual baseline subtraction using the spot tool to quantify with puncta specifications as x,y diameter of 0.18μm and z projection of 0.36μm (*68*). All images were visually inspected in 3D using the spot tool function to verify quantification.

### Pairwise and multiple sequence alignment and comparison

Multiple sequence alignment was performed on GGNBP2 (NP_079111.1, *Homo sapiens*) and its orthologues in *Mus musculus* (NP_001351040.1), *Danio rerio* (NP_001073430.2) and *Drosophila melanogaster* (NP_649570.1) (*69*). Pairwise sequence alignment and comparison was made between GGNBP2 (*Homo sapiens*) and each orthologue using EMBOSS needle (EMBL) to assess sequence identity and similarity (*15*).

### Fly RNA isolation, library preparation and sequencing

Three biological replicates from five genotypes (*Control (w^1118^)*, *Actin-Gal4, dGgnbp2^null^*, d*Ggnbp2 ^EPgy2^ (insertion only)*, and d*Ggnbp2^OE-EPgy2^* (*Actin::dGgnbp2^EPgy2^*)) were collected from the third instar larval nervous system (third instar larval brain, VNC and imaginal discs). Samples were dissected in iced PBS-DEPC water using clean tools (RNAse Zap (Sigma R2020) prior to use) before being spun (5000rpm 4⁰C for 3 minutes) and homogenised (70 seconds with a pestle) in 300 ul trizol (Invitrogen 15596-026). RNA was extracted using the Zymo Direct-zol RNA Microprep Kit (Zymo Research) as per manufacturer’s instructions and stored at -80C until library preparation.

The RNA Seq library was generated using Illumina Tru-Seq stranded mRNA kit (Illumina 20020594) as per manufacturer’s instructions. Average library fragments ranged from 316-323 bp and QC was performed using the electropherogram from an Agilent BioAnalzer. RIN scores were calculated but not used for QC due to denaturation of the 28S rRNA subunit in insect RNA. Paired-end sequencing was carried out using the Illumina NovaSeq 6000.

### RNA-seq QC and data analysis

Raw sequencing data was aligned to the *Drosophila* melanogaster genome (Release 6 + ISO1 MT/dm6) in Galaxy. The 15 samples (3 replicates for 5 different *Drosophila* lines (each sample included DNA from 30 flies)) had, on average, 55.6 (standard deviation (SD) 6.0) million reads mapped to the 15,313 genes in *Drosophila* genome. They had very similar distributions of counts per gene and total number of counts per sample. Gene with less than 10 reads across all samples (n=2334 removed) leaving 12,979 genes ‘with non-zero total reads’) for the subsequent analyses.

A ‘line-check’ was performed using the *dGgnbp2* expression levels (*CG2182*) which confirmed the genetic transformations and sequencing results were consistent with expectation (Figure 6). Briefly, there was a 7.5-fold increase in expression in the *dGgnbp2^OE-EPgy2^* vs the control *w^1118^* (58,690 ±SD 2266 vs 8206 ±SD 766 reads, respectively). While there were only 25±SD 7 reads per sample in the *dGgnbp2^null^* to demonstrate near-complete knockout. For samples with insertion of the UAS element only (d*Ggnbp2*^EPgy2^ (*Actin::Ggnbp2*^EPgy2^)), *Ggnbp2* expression was less than half (42%) of the control *w^1118^*.

To investigate if samples from the same lines clustered together, a matrix of pairwise sample distances was derived. This was carried out using using variance stabilizing algorithms on the count matrix, transformed from raw reads (*70-72*). A principal components analysis of the sample distance matrix was then used to formally identify an outlier sample. Examination of the pair-wise sample distance matrix identified one sample from *dGgnbp2^OE-EPgy2^* (*Actin::dGgnbp2*^EPgy2^) that did not cluster with the other replicates from the line indicating a potential technical problem for this sample which was excluded from downstream analyses. After exclusion of this sample, the first principal component of the distance matrix was associated with line and the second with *dGgnbp2* expression.

### Statistical Analysis of gene expression data

Following QC, a differential expression analysis of the processed count data was carried out using DESeq2 (*73*) (http://www.bioconductor.org/packages/release/bioc/html/DESeq2.html) in R. Briefly, DESeq2 assumes that the read counts for genes follow a negative binomial distribution and estimates a gene specific dispersion parameter from the input data. Differences in gene expression between pairs of lines are expressed as log2 fold changes calculated using the Approximate Posterior Estimation for generalised linear model (*74*). Consistent with DESeq2 best practice guidelines the ‘apeglm’ option was implemented in the shrinkage function ‘lfcShrink’ (*73*).

The five *Drosophila* lines (*Control (w^1118^)*, *Actin-Gal4, dGgnbp2^null^*, *dGgnbp2^EPgy2^ (insertion only)*, and *dGgnbp2^OE-EPgy2^* (*Actin::dGgnbp2^EPgy2^*)) were analysed jointly in DESeq2. Four contrast effects were defined and compared to the *Control (w^1118^*) line. This included a technical effect (affecting gene expression in both *dGgnbp2^EPgy2^* and *Actin-Gal4* line) and three biological effects (*dGgnbp2* over-expression in *dGgnbp2^OE-EPgy2^*line, reduced *dGgnbp2* expression in *dGgnbp2^EPgy2^* line, and loss of *dGgnbp2* expression in *dGgnbp2^null^* line). The log2 transformed fold change was estimated for each gene, for each effect. To correct for multiple testing the p-value was adjusted using the Benjamini and Hochberg method was to adjust the p-value (*73*). Differentially expressed genes were selected based on an adjusted p-value < 0.05 were selected. Any gene significantly associated with the *Actin-Gal4* contrast was excluded from the list of genes found as significantly associated in the *dGgnbp2^OE-EPgy2^* contrast.

Specifically, the Rcode used was:

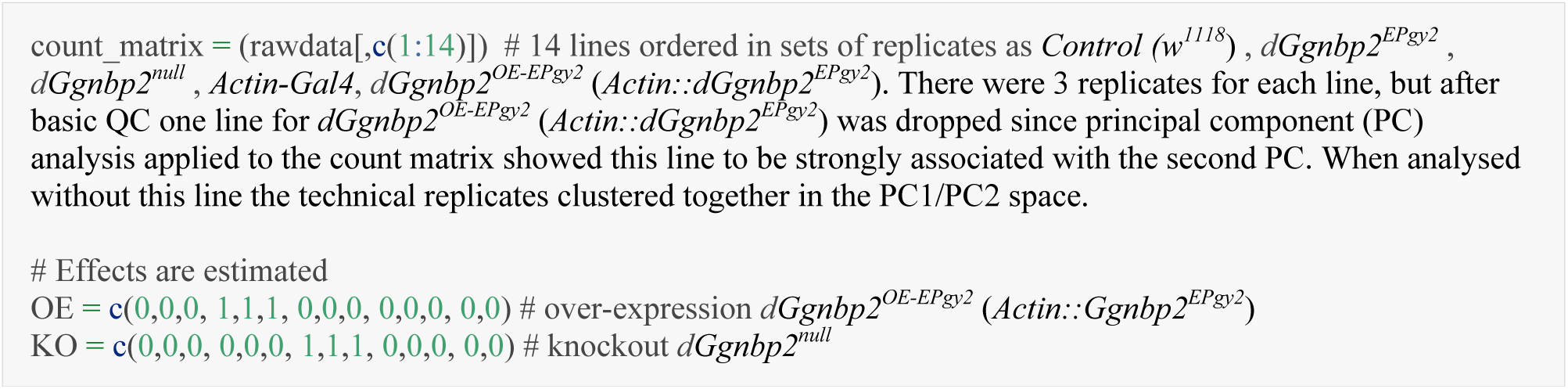
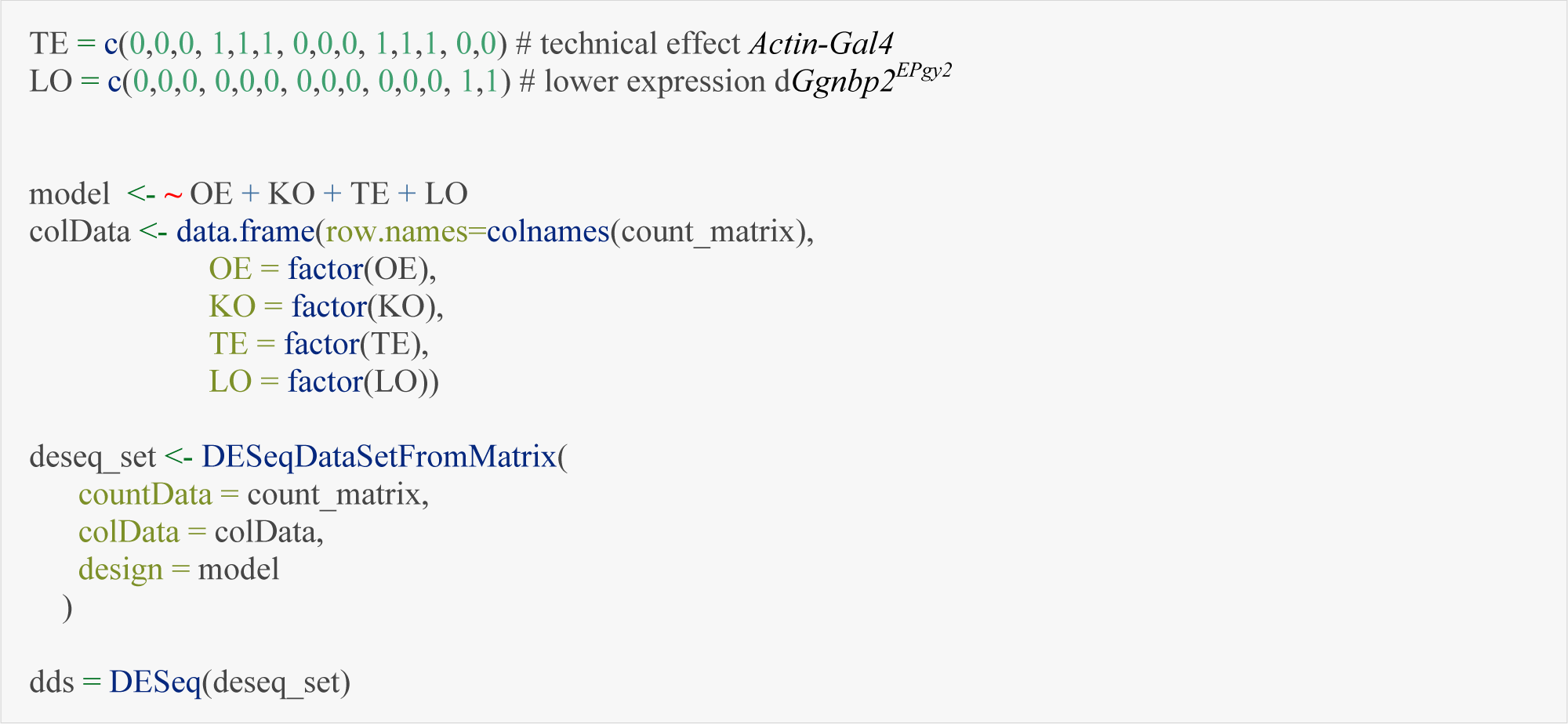

### Differentially expressed genes affected by *dGgnbp2* overexpression and *dGgnbp2* knockout

At the adjusted p-value threshold, over 10% of all genes were differentially expressed in each line. A reasonable expectation is that genes affected by *dGgnbp2* should have inverse relationships in OE lines and KO lines. A small percentage (83 genes) were significantly associated with both OE and KO effect but with signs in opposite directions. This included *CG2182 or dGgnbp2* (*GGNBP2 **Drosophila melanogaster*** ortholog).

| Effect estimated in the linear model | Number up-regulated | Number down-regulated | Total |
| --- | --- | --- | --- |
| OE-EPgy2 (*dGgnbp2* overexpression) | 1080 | 1230 | 2310 |
| OE-EPgy2 but not TE* | 703 | 821 | 1524 |
| KO (*dGgnbp2<sup>null</sup>*) | 2407 | 2365 | 4772 |
| LO (*dGgnbp<sup>EPgy2</sup>*) | 1713 | 1719 | 3432 |
| TE (technical effect of GAL4 insertion) | 1749 | 1554 | 3303 |

\*786 genes in the TE list (indicating that the expression in TE\_L differed from both WT\_L and OE\_L) were excluded from the OE list

### Functional enrichment analysis using g:Profiler of differentially expressed genes

g:Profiler (version e109_eg56_p17_1d3191d) was used for functional enrichment analysis with g:SCS multiple testing correction method with a significance threshold of 0.05 (*33*).

### Negative Geotaxis

Flies were grouped into cohorts of 10 from three independent crosses (three biological replicates). Datapoints represent the average of three technical replicates that were averaged at each time point, per genotype. A vial was used in the assay with an 8cm line to perform the assay. Flies were subjected to 4 ‘taps’ and given 10 seconds to reach the line. The number of flies that passed the line divided by the total number of flies was defined as performance index (PI). This was repeated three times with a one-minute rest separating each trial.

### Quantification of autophagy and PI3K markers at the third instar larval NMJ

Quantification of corrected fluorescence intensity using integrated density (arbitrary units A.U.) of *pm-Atg8::mCherry*, *Ref(2)P.GFP*, PIP_2_ and *GFP-myc-2xFYVE* in 1b motor neurons was obtained using HRP as a guide for boutons in ImageJ. Ten boutons from 1b were averaged. Three non-synaptic areas of similar surface area were averaged and subtracted to account from background. For the punctate labelling of *Ref(2)P.GFP*, PIP_2_ and *GFP-myc-2xFYVE* puncta density (number of puncta per μm^2^) and puncta corrected fluorescence intensity was measured.

### Statistical Analyses of fly motor neuron morphology

Statistical analyses for fly motor neuron morphology were performed using Prism 7.0 software (GraphPad). D’Agostino & Pearson omnibus normality tests were conducted to determine whether each fly dataset followed a Gaussian (normal) distribution. The significance for normality test was set at p<0.05 (where statistical significance equates to a dataset that does not follow a normal distribution). Normally distributed datasets were subject to a t-test or parametric ANOVA test and a Tukey’s multiple comparison tests. Datasets containing non-Gaussian distributions were subjected to a Mann-Whitney test or Kruskal-Wallis test (non-parametric ANOVA) and a Dunn’s multiple comparisons test. For categorical data we conducted Fischer’s exact test. Negative geotaxis was subjected to a Two-way ANOVA with a Tukey’s or Sidak multiple comparison test. Statistical significance in figures are depicted with asterisks as follows: n.s p>0.05, *\*p*<0.05, \**\*p*<0.01, \*\**\*p*<0.001 and \*\*\**\*p*<0.0001.

## Acknowledgments

We would like to thank Donna Denton for kindly sharing several fly transgenic lines to assess autophagy, Adam Walker and Stefan Thor for providing feedback on the manuscript.

## Funding

S.K.K. was supported by an Australian Postgraduate Award (Research Training Scheme) and UQ Research Fellowship. This study was supported by an Australian National Health and Medical Research Council (NHMRC) grant (1173790, 113400), a Scott Sullivan MND Fellowship from the MND and Me Foundation, Motor Neuron Disease Research Australia (to F.C.G). B.M.C. is supported by NHMRC Senior Research Fellowship and Investigator Grant (1136021, 2016410).

### Author contributions

S.K.K., S.S.M and N.R.W. were responsible for conceptualization, experimental design and manuscript preparation. F.C.G. also contributed significantly to manuscript preparation. S.K.K performed, collected and analysed data for fly motor neuron and nuclear morphology, autophagy and signalling markers. S.K.K performed GO analysis on RNA-seq data. S.K.K., N.C. and A.K. performed, collected and analysed data for fly behavioural experiments. N.C. performed fly embryo microinjections for CRISPR and transgenic lines and prepared fly RNA for RNA-seq. T.L., N.R.W. and T.L. designed and analysed the RNA-seq QC and DE analysis. A.K. performed, collected and analysed data for fly Ggnbp2 localisation and GluRIIA in motor neurons. N.R.W, X.Q., A.F.M., and F.C.G. conceived and performed the post-GWAS human analyses. B.C. generated and aligned the human and fly Alphafold2 structures.

## Data and materials availability

All data are available in the main text or the supplementary materials.

## Table of contents for supplementary materials

## Supplementary Text

Figure S1. Multiple sequence alignment of *GGNBP2*

Figure S2. Sequence alignment of *H. sapiens* and *D. melanogaster* GGNBP2 proteins

Figure S3. *dGgnbp2^null^* motor neuron morphology defects are presynaptic

Figure S4. Overexpression of *dGgnbp2* alters motor neuron morphology

Figure S5. Locomotion defects in females overexpressing or lacking *dGgnbp2*

Figure S6. *hGGNBP2* rescues *dGgnbp2* motor neuron morphology and locomotion defects

Figure S7. Cytoplasmic dGgnbp2 is sufficient to rescue *dGgnbp2^null^* motor neuron defects

Figure S8. Differential gene expression upon loss and overexpression of *dGgnbp2*

Figure S9. *dGgnbp2* is linked to *ik2* and regulates lipid phosphorylation

Table S1. Conditional and Joint analysis (GCTA-COJO)

Table S2. Gene-based test of ALS GWAS using FastBAT and mBAT Table S3. SMR results from blood (eQTLgen)

Table S4. Rare variant burden in ALS cases (n=4633) vs controls (n=1832)

Table S5. SMR results from brain (metabrain)

Table S6a. Genes that change in opposite directions between OE and KO

Table S6b. Genes that change in the same direction between OE and KO

Table S6c. Full RNA-seq results

## Figures and Tables

**Fig. S1.**
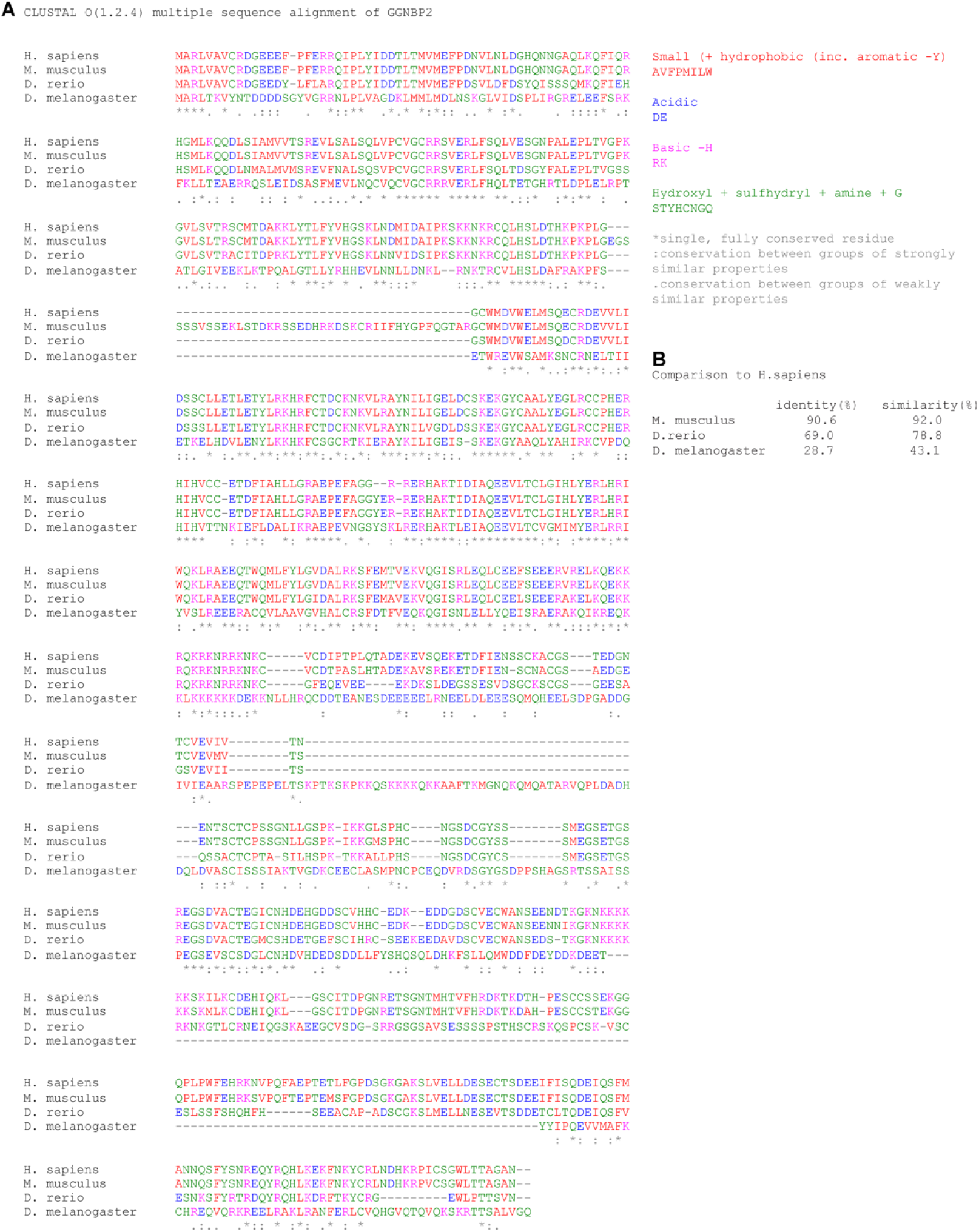
Multiple sequence alignment of *GGNBP2*. (A) Multiple sequence alignment was performed on GGNBP2 (NP_079111.1, *Homo sapiens*) and its orthologues in *Mus musculus* (NP_001351040.1), *Danio rerio* (NP_001073430.2) and *Drosophila melanogaster* (NP_649570.1). (B) Pairwise sequence alignment and comparison were made between GGNBP2 (*Homo sapiens*) and each orthologue using EMBOSS needle (EMBL) to assess sequence identity and similarity.

**Fig. S2.**
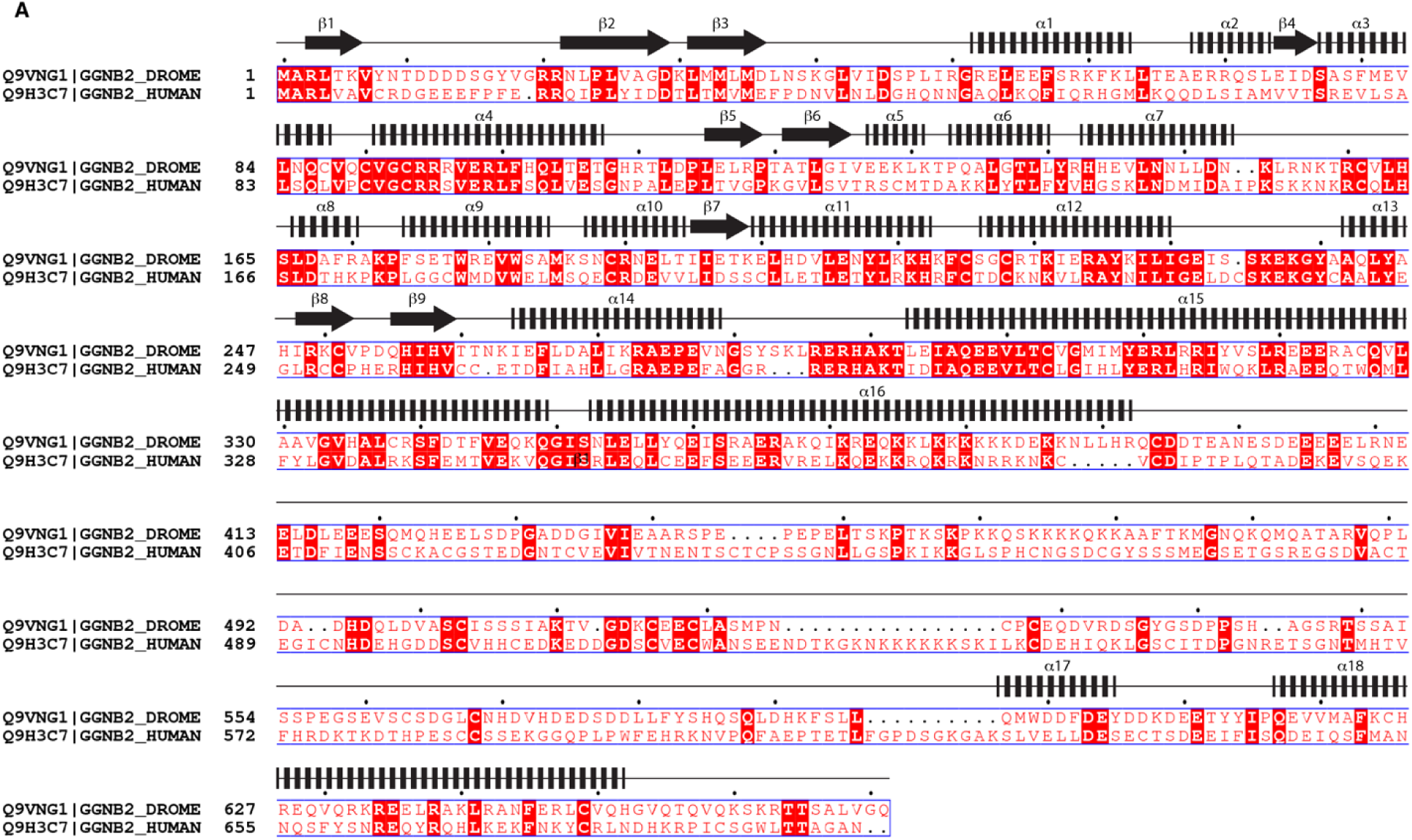
Sequence alignment of *H. sapiens* and *D. melanogaster* GGNBP2 proteins. (A) Sequence alignment of the fly and human GGNBP2 proteins with the predicted secondary structure of fly dGgnbp2 (CG2182) indicated. Conserved amino acids are in red.

**Fig. S3.**
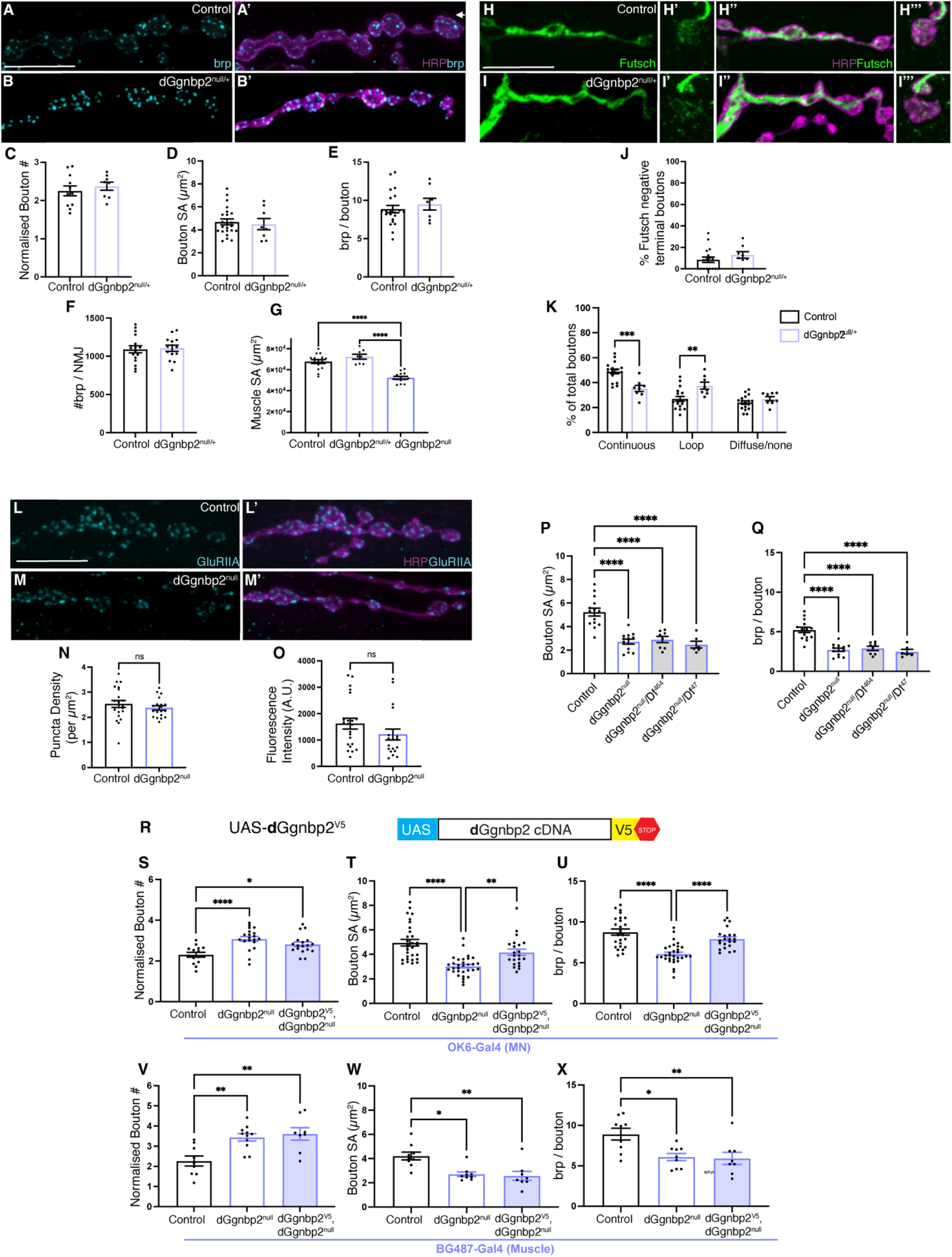
*dGgnbp2^null^* motor neuron morphology defects are presynaptic. (A-A’) Control (*w^1118^*) MN6/71b labelled with nc82 antibody (cyan, A) to mark brp positive active zones and merged with HRP (magenta, A’) which labels the axonal membrane. (B-B’) d*Ggnbp2^null/+^* MN6/7 1b with the same labelling scheme as A. (C-G) Quantification of bouton number normalised to muscle 6/7 surface area (1 x 10^3^μm^2^) (C) bouton size (D) brp puncta/bouton (E) number of brp puncta/neuromuscular junction (F) and muscle 6/7 surface area (1 x 104μm^2^) (G) n=11-22. (H-I) Futsch labelling of Control (*w^1118^*) MN6/71b labelled with antibody to futsch (green, H-H’) and merged with HRP (magenta, H’’-H’’’). High magnification of a terminal bouton showing a futsch loop (H’, H’’’). *dGgnbp2^null/+^* motor neuron with the same labelling scheme as in H (I-I’’’). (J-K) Quantification of futsch in terminal boutons (J) and the different states of futsch labelling in control and *dGgnbp2^null/+^* boutons. n=8-9 (K). (L-L’) The motor neuron terminal of control (*w^1118^*) at muscle 6/7 labelled by GluRIIA (cyan, L) and merged with HRP to label the axonal membrane (magenta, L’) in 3rd instar *Drosophila* larvae. *dGgnbp2^null^* motor neuron with the same labelling scheme as in N (M- M’). (N-O) Quantification of GluRIIA puncta density per bouton (μm2) (N) and fluorescence intensity (A.U.) per bouton (O) n=18-20. (P-Q) Quantification in MN6/71b for bouton surface area (μm2) (P) and number of brp puncta per bouton (Q) for *Control^w1118^*, *dGgnbp2^null^*, trans heterozygotes of chromosomal deficiency lines (*Df464*, *Df47*) and *dGgnbp2^null^* n= 6-13. (R) Schematic for *UAS-dGgnbp2^V5^* construct. (S-U) Quantification of motor neuron phenotypes as in C-E following rescue with *dGgnbp2* in motor neurons (*OK6-Gal4*) n=14-27. (V-X) Quantification of motor neuron phenotypes as in C-E following rescue with *dGgnbp2* in muscle (*BG487-Gal4*) n=8-11. All quantitative analyses were performed on compressed Z stacks of spinning disk microscopy and analysed using ImageJ. Each data point=average of 10 boutons. Depending on distribution of data an unpaired t test (C-F, N) or Mann-Whitney test (O) or an Ordinary one-way ANOVA with Tukey’s multiple comparison test (G, J-K, P-Q, S, U-V, X) or Kruskal-Wallis test with Dunn’s multiple comparison test (T,W). n.s *p*>0.05, \**p*<0.05, \*\*\**p*<0.001 and \*\*\*\**p*<0.0001. White arrows show MN6/71b. Scale bars denote 10um.

**Fig. S4.**
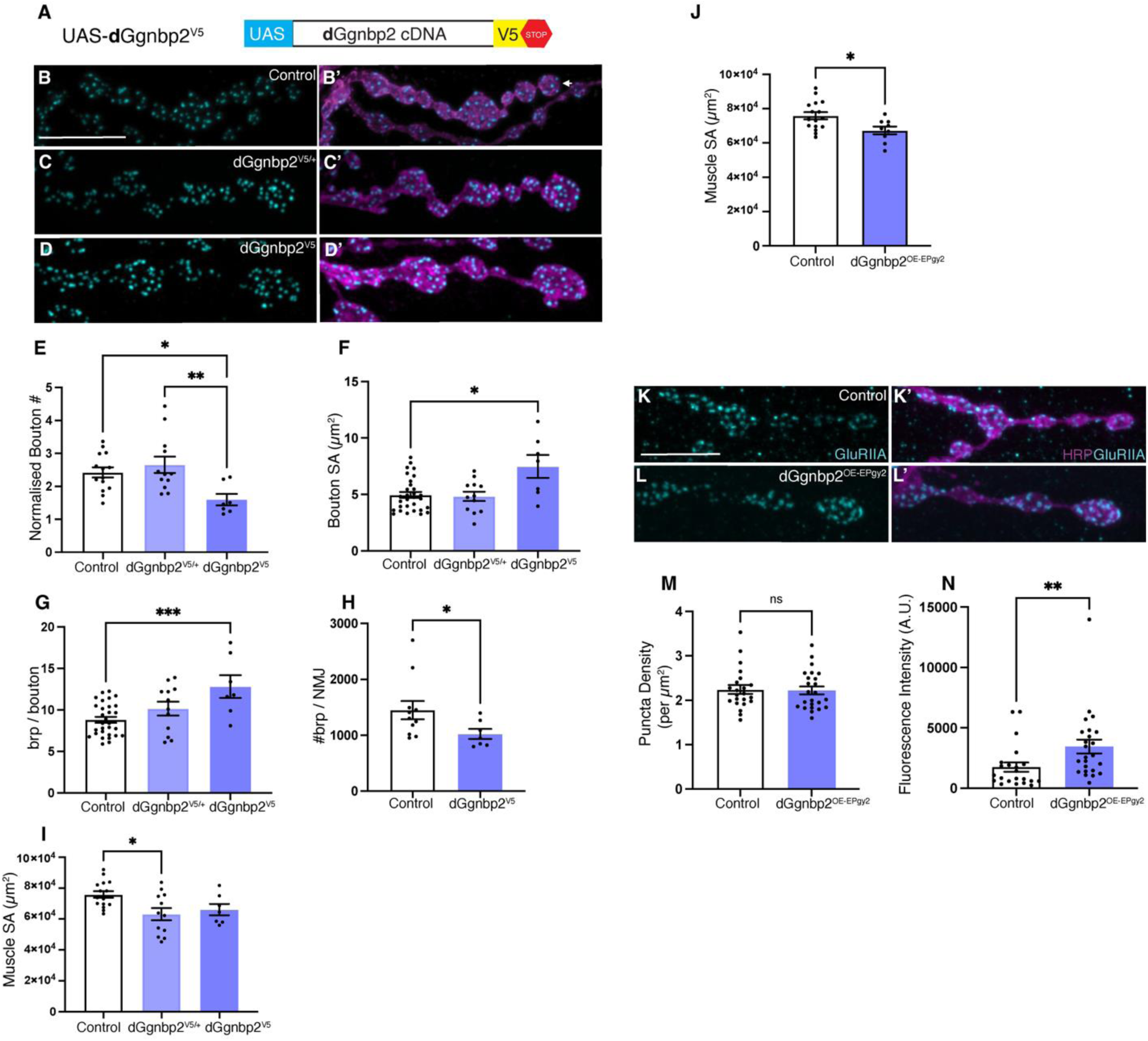
Overexpression of *dGgnbp2* alters motor neuron morphology. (A) Schematic for *UAS-dGgnbp2^V5^* construct. (B-B’) Control (*OK6-Gal4*) MN6/71b labelled with nc82 antibody (cyan, B) to mark brp positive active zones and merged with HRP (magenta, B’) which labels the axonal membrane. (C-C’) *OK6-Gal4* driving expression of *UAS- dGgnbp2^V5^/+* MN6/7 1b with the same labelling scheme as G. (D-D’) *OK6-Gal4* driving expression of *UAS-dGgnbp2^V5^* MN6/7 1b with the same labelling scheme as B. (E-I) Quantification of bouton number normalised to muscle 6/7 surface area (1 x 10^3^μm2) (E) bouton size (F) brp puncta/bouton (G) number of brp puncta/neuromuscular junction (H) and muscle 6/7 surface area (1 x 104μm2) (I) n=7-25. (J) Quantification of muscle 6/7 surface area (1 x 10^4^μm^2^) in *Control^OK6-Gal4^* and *dGgnbp2^OE-EPgy2^* n=9-16. (K-K’) The motor neuron terminal of control (*w^1118^*) at muscle 6/7 labelled by GluRIIA (cyan, K) and merged with HRP to label the axonal membrane (magenta, K’) in 3rd instar *Drosophila* larvae. *dGgnbp2^OE-EPgy2^*motor neuron with the same labelling scheme as in N (L-L’). (M-N) Quantification of GluRIIA puncta density per bouton (μm2) (M) and fluorescence intensity (A.U.) per bouton (N) n=18-20. All quantitative analyses were performed on compressed Z stacks of spinning disk microscopy and analysed using ImageJ. Each data point=average of 10 boutons. Depending on distribution of data an unpaired t test (J) or Mann-Whitney test (H, M-N) or an Ordinary one-way ANOVA with Tukey’s multiple comparison test (E, G, I) or Kruskal-Wallis test with Dunn’s multiple comparison test (F). n.s *p*>0.05, \**p*<0.05, \*\*\**p*<0.001 and \*\*\*\**p*<0.0001. Scale bars denote 10um.

**Fig. S5.**
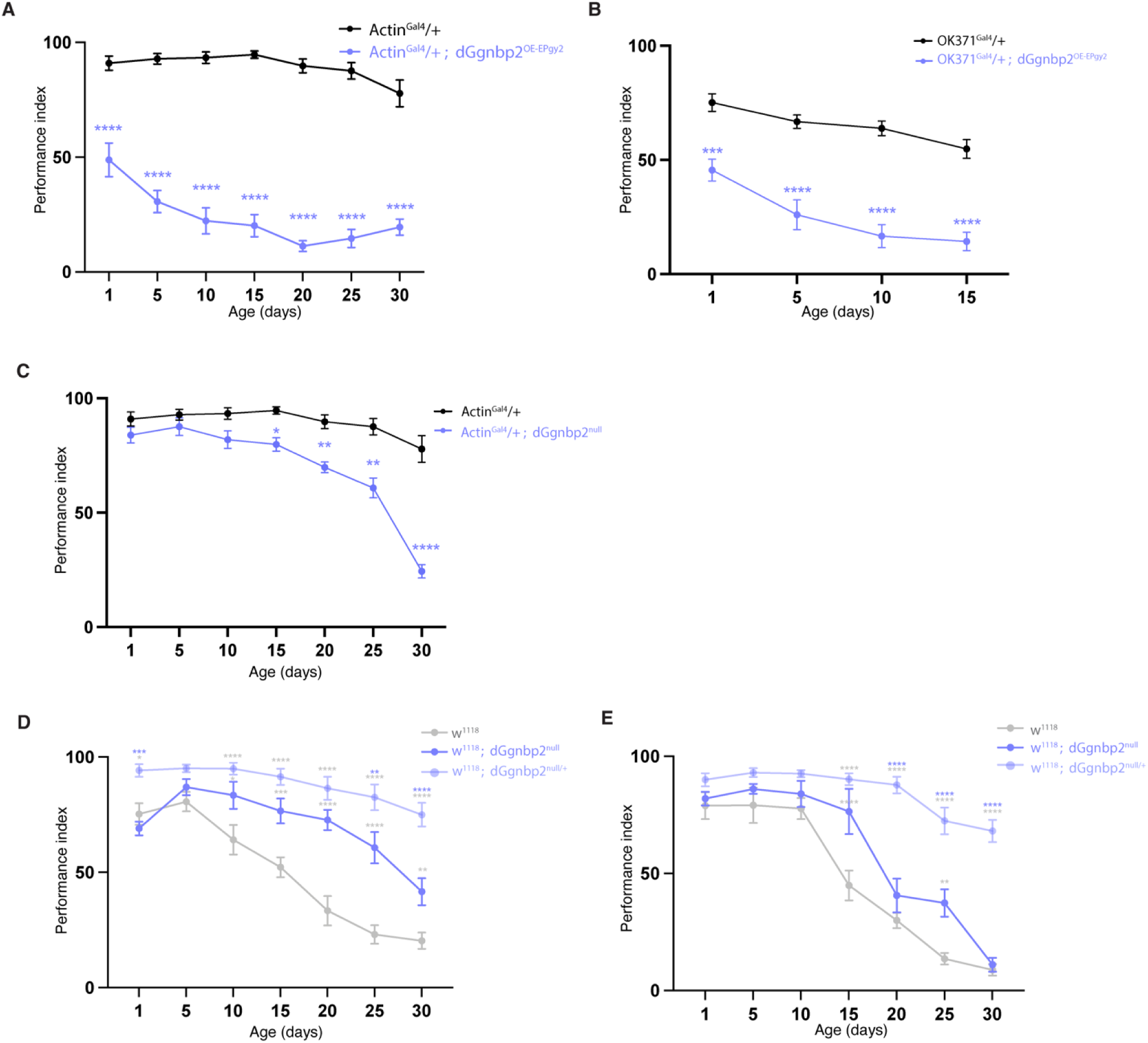
Locomotion defects in females overexpressing or lacking *dGgnbp2*. (A) *Control^Actin-Gal4^* (black) and *dGgnbp2^OE-EPgy2^*(purple) females were assessed for performance index (PI) over 30 days. (B) *Control^OK371-Gal4^* (black) and *dGgnbp2^OE-EPgy2^* (purple) females overexpressing *dGgnbp2* in motor neurons. (C) *Control^Actin-Gal4^*(black) and *dGgnbp2^null^*(purple). (D) *Control^w1118^* males and females (E) were compared to *dGgnbp2^null^* and *dGgnbp2^null/+^* on a *w^1118^* background. Flies were grouped into cohorts of 10 flies and generated from three independent crosses (three biological replicates). Data represents the average of three technical replicates that were averaged at each time point per genotype. n=7-9 cohorts of 10 flies. Data are represented as mean ± SEM. A Two-way ANOVA was performed with a Tukey’s or Sidak multiple comparison test. n.s *p*>0.05, \**p*<0.05, \*\*\**p*<0.001 and \*\*\*\**p*<0.0001.

**Fig. S6.**
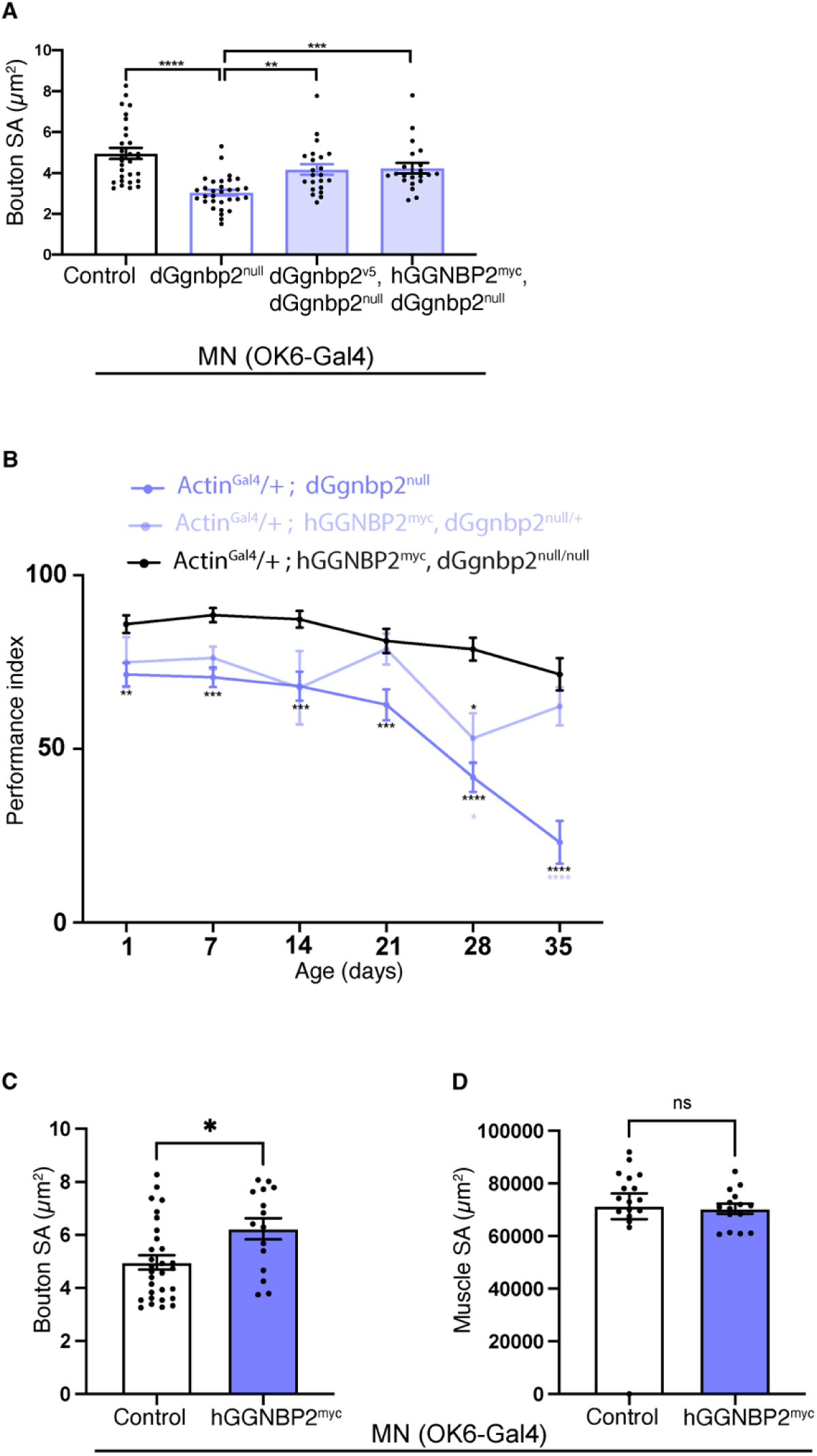
h*GGNBP2* rescues *dGgnbp2* motor neuron morphology and locomotion defects. (A) Rescue of *dGgnbp2^null^* motor neuron phenotypes with fly and human *GGNBP2*. Quantification of bouton size (A) in the four different genotypes: Control (*OK6-Gal4*), *dGgnbp2^null^*, *OK6-Gal4*/ *dGgnbp2^V5^*; *dGgnbp2^null^, OK6-Gal4*/ *hGGNBP2^myc^*; *dGgnbp2^null^*. n=21-31. (B) Rescue of *dGgnbp2^null^*locomotion phenotypes with *hGGNBP2* in females that were assessed for PI over 35 days. Flies were grouped into cohorts of 10 flies and generated from three independent crosses (three biological replicates). Data represents the average of three technical replicates that were averaged at each time point per genotype. n=7-9 cohorts of 10 flies. Data are represented as mean ± SEM. A Two-way ANOVA was performed with a Tukey’s or Sidak multiple comparison test (C-D) Quantification of bouton size (C) and muscle 6/7 surface area (μm^2^) (D) in *Control* (*OK6-Gal4*) and *hGGNBP2^myc^* (*OK6-Gal4:: hGGNBP2^myc^*) n=16-30. All quantitative analyses were performed on compressed Z stacks of spinning disk microscopy and analysed using ImageJ. Each data point=average of 10 boutons. A Kruskal-Wallis test with Dunn’s multiple comparison (A) or unpaired t-test (C-D) was performed. n.s *p*>0.05, \**p*<0.05, \*\*\**p*<0.001 and \*\*\*\**p*<0.0001.

**Fig. S7.**
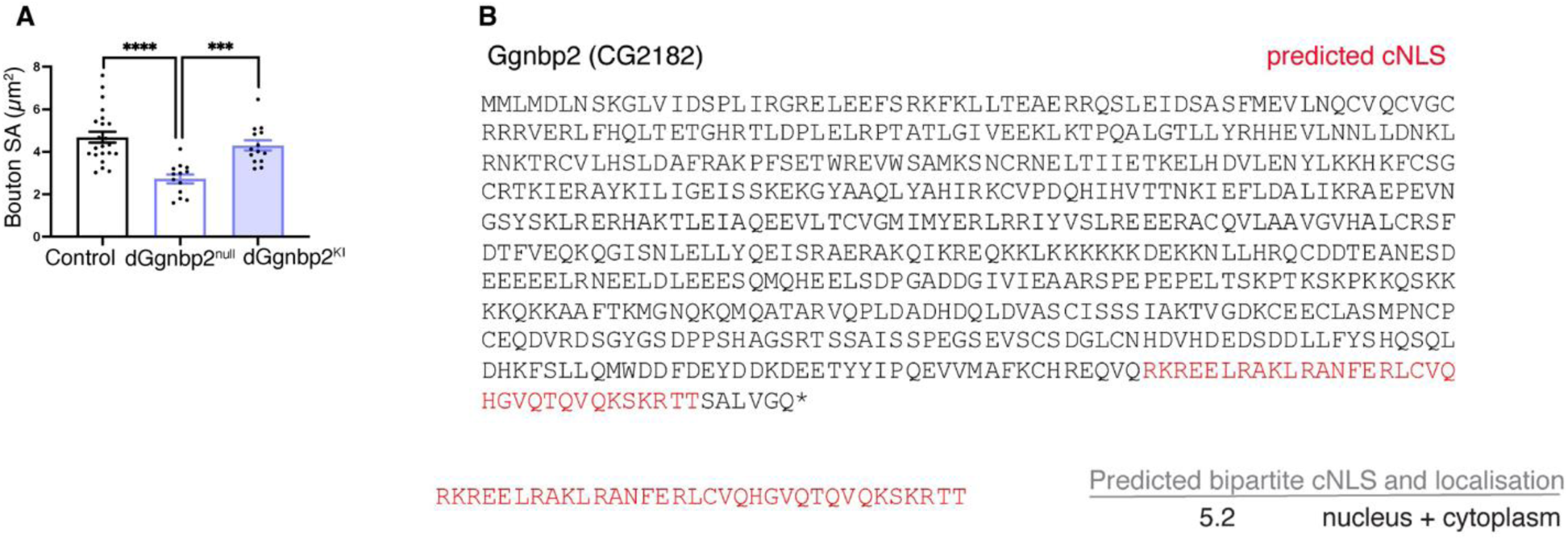
Cytoplasmic *dGgnbp2* is sufficient to rescue *dGgnbp2^null^* motor neuron defects. (A) Quantification of MN6/71b bouton size in *Control^w1118^*, *dGgnbp2^null^*and *dGgnbp2^KI-V5^*. n=12-20. All quantitative analyses were performed on compressed Z stacks of spinning disk microscopy and analysed using ImageJ. Each data point=average of 10 boutons. Data are represented as mean ± SEM. An Ordinary one-way ANOVA with Tukey’s multiple comparison test (A). n.s *p*>0.05, \**p*<0.05, \*\*\**p*<0.001 and \*\*\*\**p*<0.0001. (B) Protein sequence of fly Ggnbp2 (CG2182) and the result of cNLS prediction software. Predicted cNLS (red).

**Fig. S8.**
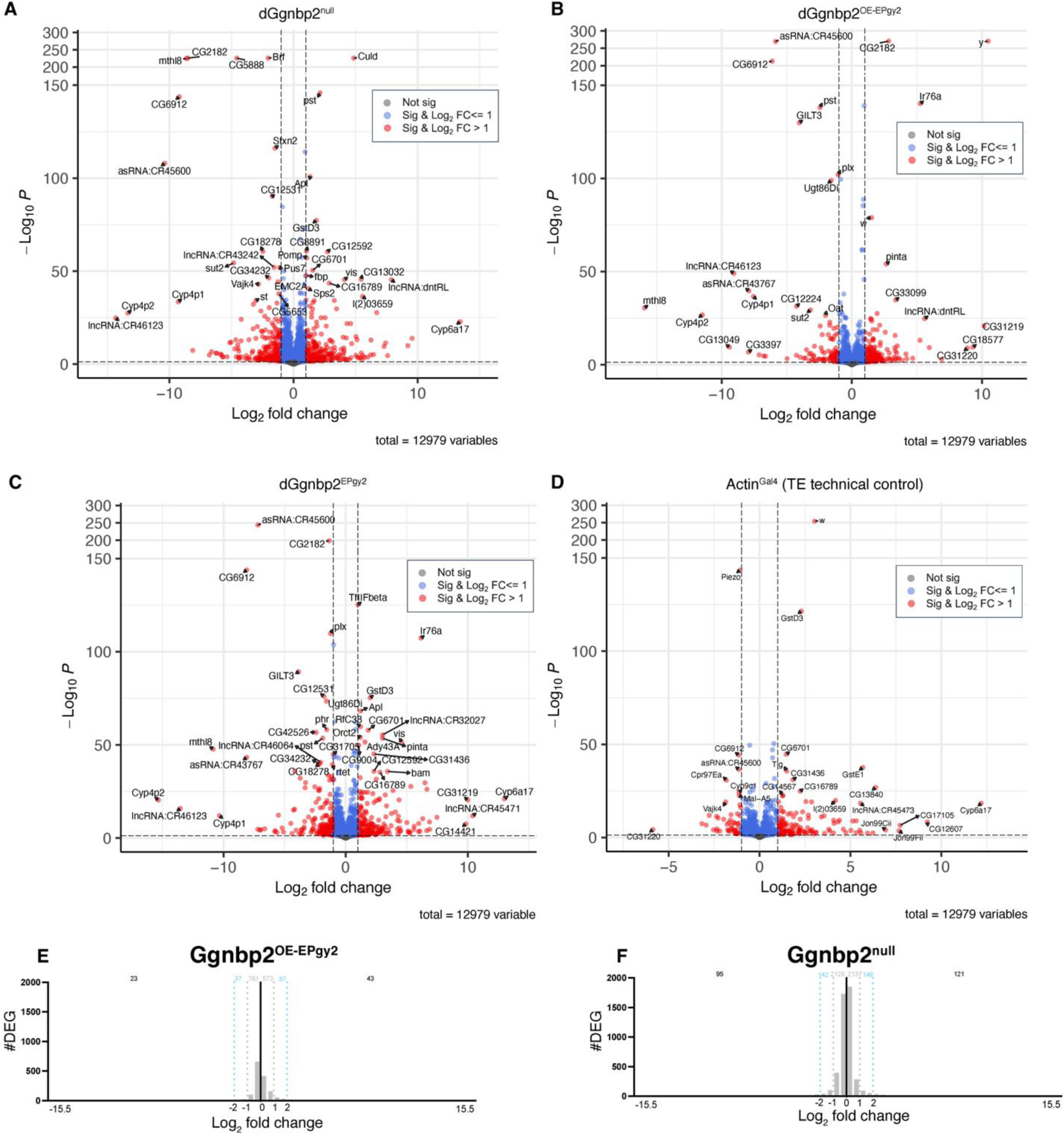
Differential gene expression upon loss and overexpression of *dGgnbp2*. (A-D) Volcano plots of the RNA-seq data from the 4 different genotypes: *dGgnbp2^null^*(A), *dGgnbp2^OE-EPgy2^* (B), *dGgnbp2* (C) and *Actin-Gal4* (D). Log_2_ fold change in gene expression is shown on the x axis plotted against -Log_10_ p value on the y axis. The y axis range from 150 to 300 has been shortened to enable better visualization of the genes at the bottom of the graph. Genes are represented by a circle colored according to their significance and magnitude of change in gene expression (not significant – grey, significant and a log_2_ fold change less than 1 – blue, significant with a log_2_ fold change greater than 1 – red). Gene names are indicated when known. (E-F) Number of differentially expressed genes (DEG) and their log_2_ fold change in *dGgnbp2^OE-EPgy2^* and *dGgnbp2^null^*.

**Fig. S9.**
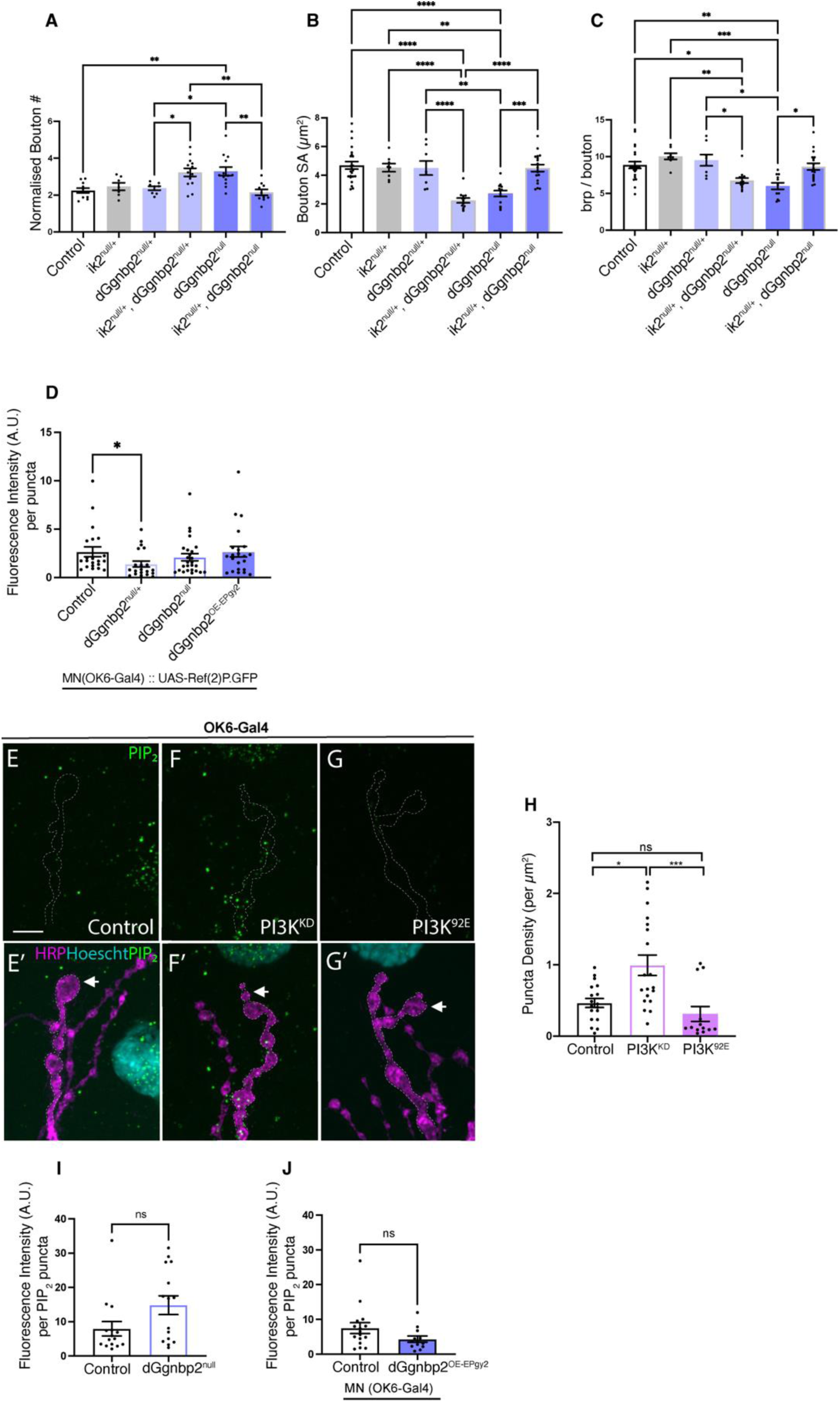
*dGgnbp2* is linked to *ik2* and regulates lipid phosphorylation. (A-C) Genetic interactions between *dGgnbp2* and *ik2* in motor neurons. Quantification of (A) bouton number/1 x 10^4^ μm^2^ of muscle (B) bouton size and (C) number of brp puncta per bouton in different *dGgnbp2* and *ik2* genotypes (D) Quantification of Ref(2)P.GFP puncta fluorescence intensity (A.U.) in different *dGgnbp2* genotypes all expressing *OK6-Gal4::UAS-Ref(2)P.GFP.* (G-I’) MN6/71b labelled with nuclear marker Hoescht (cyan), HRP (E-G) and motor neuron terminal outlined is overlayed (E-G) on images for an antibody against PIP_2_ (green puncta) in Control (*OK6-Gal4*, E-E’), *PI3K^KD^* (F-F’), *PI3K^92E^* (G-G’). (H) Quantification of PIP_2_ puncta density (per μm^2^). n=13-19. (I) Quantification of PIP_2_ puncta fluorescence intensity (A.U.) in *Control^w1118^*and *dGgnbp2^null^* using an antibody to PIP_2_. (J) Quantification of PIP_2_ puncta fluorescence intensity (A.U.) in *Control^OK6-Gal4^*and *dGgnbp2^OE-EPgy2^*. All quantitative analyses were performed on compressed Z stacks of spinning disk microscopy and analysed using ImageJ. Each data point=average of 10 boutons. Data are represented as mean ± SEM. Depending on distribution of data an unpaired t test (J) or Mann-Whitney test (I) or an Ordinary one-way ANOVA with Tukey’s multiple comparison test (A-B) or Kruskal-Wallis test with Dunn’s multiple comparison test (C-D, H). n.s *p*>0.05, \**p*<0.05, \*\*\**p*<0.001 and \*\*\*\**p*<0.0001.

